# E4F1 COORDINATES PYRUVATE METABOLISM AND THE ACTIVITY OF THE ELONGATOR COMPLEX TO ENSURE TRANSLATION FIDELITY DURING BRAIN DEVELOPMENT

**DOI:** 10.1101/2022.12.19.521032

**Authors:** Di Michele Michela, Attina Aurore, Roux Pierre-François, Tabet Imène, Laguesse Sophie, Florido Javier, Houdeville Morane, Choquet Armelle, Encislai Betty, Arena Giuseppe, De Blasio Carlo, Wendling Olivia, Frenois Francois-Xavier, Papon Laura, Stuani Lucille, Fuentes Maryse, Jahanault-Tagliani Céline, Rousseau Mélanie, Guégan Justine, Buscail Yoan, Dupré Pierrick, Michaud Henri-Alexandre, Rodier Geneviève, Bellvert Floriant, Kulyk Hanna, Ferraro Peyret Carole, Mathieu Hugo, Close Pierre, Rapino Francesca, Chaveroux Cédric, Pirot Nelly, Rubio Lucie, Torro Adeline, Sorg Tania, Ango Fabrice, Hirtz Christophe, Compan Vincent, Lebigot Elise, Legati Andrea, Ghezzi Daniele, Nguyen Laurent, David Alexandre, Sardet Claude, Lacroix Matthieu, Le Cam Laurent

## Abstract

Pyruvate metabolism defects lead to severe neuropathies such as the Leigh syndrome (LS) but the molecular mechanisms underlying neuronal cell death remain poorly understood. Here, we unravel a connection between pyruvate metabolism and the regulation of the epitranscriptome that is relevant to LS pathogenesis. We identified the transcription factor E4F1 as a key coordinator of AcetylCoenzyme A (AcCoA) production by the pyruvate dehydrogenase complex (PDC) and its utilization as an essential co-factor by the Elongator complex to acetylate tRNAs at the wobble position uridine 34 (U_34_). E4F1-mediated direct transcriptional regulation of *Dlat* and *Elp3*, two genes encoding key subunits of the PDC and of the Elongator complex, respectively, ensured proper translation fidelity and cell survival in the central nervous system (CNS) during mouse embryonic development. Furthermore, analysis of PDH-deficient cells highlighted a crosstalk linking the PDC to ELP3 expression that is perturbed in LS patients.

## INTRODUCTION

The Leigh syndrome (LS, OMIM#256000) is a severe inborn neurodegenerative encephalopathy affecting one in forty thousand newborns that stems from mitochondrial deficiencies in pyruvate metabolism or in the electron transport chain (ETC). Mutations in over ninety-five genes encoded by either the nuclear or the mitochondrial genomes have been linked to this inborn metabolic disorder, including genes encoding key components or regulators of the pyruvate dehydrogenase (PDH) complex (PDC). PDC is a mitochondrial multisubunit complex that converts pyruvate into AcetylCoenzymeA (AcCoA) to fuel the tri-carboxylic acid cycle (TCA). Most LS patients display chronic lactate acidemia and bilateral necrotizing lesions of the basal ganglia and the brain stem. Affected children also develop a broad range of motor and neurological symptoms which lead to premature death during early childhood, including muscular atonia, growth and mental retardation, ataxia and vision loss^1^. No efficient cure has been identified so far for PDC-linked mitochondrial disorders although few patients have shown a partial improvement of their clinical symptoms under a ketogenic diet, a regimen that metabolically bypasses the PDC complex, or upon its pharmacological stimulation by dichloro-acetate (DCA)^2^. Paradoxically, despite most of the genetic alterations implicated in the LS have been identified, the molecular mechanisms underlying the clinical symptoms of these children, and in particular their neurological defects, remain poorly understood.

Converging evidence suggests that deregulation of the multifunctional protein E4F1 is associated to PDH deficiency and to the LS. The E4F1 protein was originally identified as a transcriptional regulator of the E4 viral gene that is hijacked by the viral oncoprotein E1A during adenoviral infection^3^. Since, E4F1 was also characterized as an atypical E3 ligase of the tumor suppressor p53 that controls its transcriptional activities independently of degradation^4^. Beside its role in regulating the balance between cell proliferation and cell death^4–6^, E4F1 was also identified as a major regulator of the PDC complex. Thus, E4F1 directly binds to the promoter region of several genes encoding essential subunits or regulators of the PDC complex, including DLAT and DLD, the E2 and E3 core subunits of the PDC, the regulatory subunit of the PDH phosphatase complex PDPR, the pyruvate transporter of the inner mitochondrial membrane MPC1, and the SLC25A19 protein involved in mitochondrial uptake of thiamine pyrophosphate, an essential co-factor of the PDC^7–10^. A potential connection between E4F1 and the LS was identified through the clinical management of two siblings of an italian family harboring a homozygous *E4F1*^K144Q^ missense mutation who developed clinical symptoms reminiscent of those observed in LS patients, including muscular hypotonia, microcephaly, hyperlactacidemia and organic aciduria associated to impaired PDH activity in their skeletal muscles. Brain MRI-imaging revealed symmetric necrotic lesions in the subthalamic nuclei and in the substantia nigra that are common in LS patients^11^. Consistent with a pivotal role of E4F1 in pyruvate metabolism and LS pathogenesis, knock-out (KO) mice lacking E4F1 in their skeletal muscles exhibit decreased PDH activity and display phenotypes that are reminiscent of some of the clinical symptoms of LS patients, including chronic lactate acidemia and exercice intolerance^8^. However, no major neuropathological abnormalities were observed in this animal model, suggesting that the neuropathy of LS patients results from tissue specific defects in the central nervous system (CNS). Here, we generated mice lacking E4F1 in the CNS to investigate its role during neuronal development. Detailed molecular characterization of this animal model during embryonic development and of E4F1-deficient primary fibroblasts demonstrated that in addition to its role in pyruvate metabolism, E4F1 is also a key transcriptional regulator of *Elp3*, a gene encoding the catalytic subunit of the Elongator complex. This complex is an acetyl-transferase which targets a subset of tRNAs in the acceptor stem loop at the wobble position uridine 34 (U^34^) to promote a stepwise cascade of chemical modifications (methoxy-carbonyl-methylation and thiolation) to produce mature tRNAs and thereby ensure protein translation fidelity^12–14^. Our data indicate that perturbation of E4F1 activities during embryonic development resulted in severe brain defects and microcephaly associated to impaired tRNAs U_34_ modifications, translation defects and the induction of an integrated stress response (ISR) leading to neuronal cell death. Moreover, we identified a previously unknown crosstalk between the PDC and ELP3 expression that is perturbed in cells isolated from LS patients. Altogether, our data highlight the importance of the links between pyruvate metabolism and the epitranscriptome to ensure proteome homeostasis in the CNS.

## RESULTS

### E4F1 plays a critical role in neuronal development

To evaluate the roles of *E4f1* during neuronal development, we generated animals lacking E4F1 in their CNS by crossing *E4f1* constitutive and conditional KO (cKO) mice with transgenic animals expressing the CRE recombinase under the control of the *Nestin* (*Nes*) promoter that is predominantly active in all neuronal and glial progenitors from E11, and with a reporter strain allowing Cre-mediated expression of the green fluorescent protein (GFP) from the ROSA26 locus (Supplemental Fig. S1A)^1^^5–18^. In these compound mutant animals (hereafter referred to as *E4f1^(Nes)KO^)* and their control littermates (CTL), immunohistochemistry (IHC) analysis of the GFP reporter on head or whole embryo sagittal sections isolated at different developmental stages confirmed that Cre-mediated recombination was mainly restricted to the CNS and the spinal cord (Fig. 1A and Supplemental Fig. S1B). The efficiency and the specificity of Cre-mediated inactivation of *E4f1* in the CNS was confirmed at E14.5 by RT-qPCR and immunoblotting, respectively (Fig. 1B,C). *E4f1^(Nes)KO^* mice were represented at the expected mendelian ratio at E14.5, E16.5 and E18.5 and showed no obvious growth developmental abnormalities, but were absent at P1, indicating that E4F1 deficiency in the CNS leads to perinatal death (Fig. 1D and Supplemental Fig. S1C,D). Moreover, *E4f1^(Nes)KO^* E18.5 embryos exhibited microcephaly, a developmental defect also observed in some LS patients^1^ including those harboring the homozygous *E4F1^A430C^* mutation which encodes an hypomorpic allele of E4F1, as shown by our complementation assays in *E4f1^cKO^* Mouse Embryonic Fibroblasts (MEFs) (Supplemental Fig. S1E-G).

**Figure 1.**
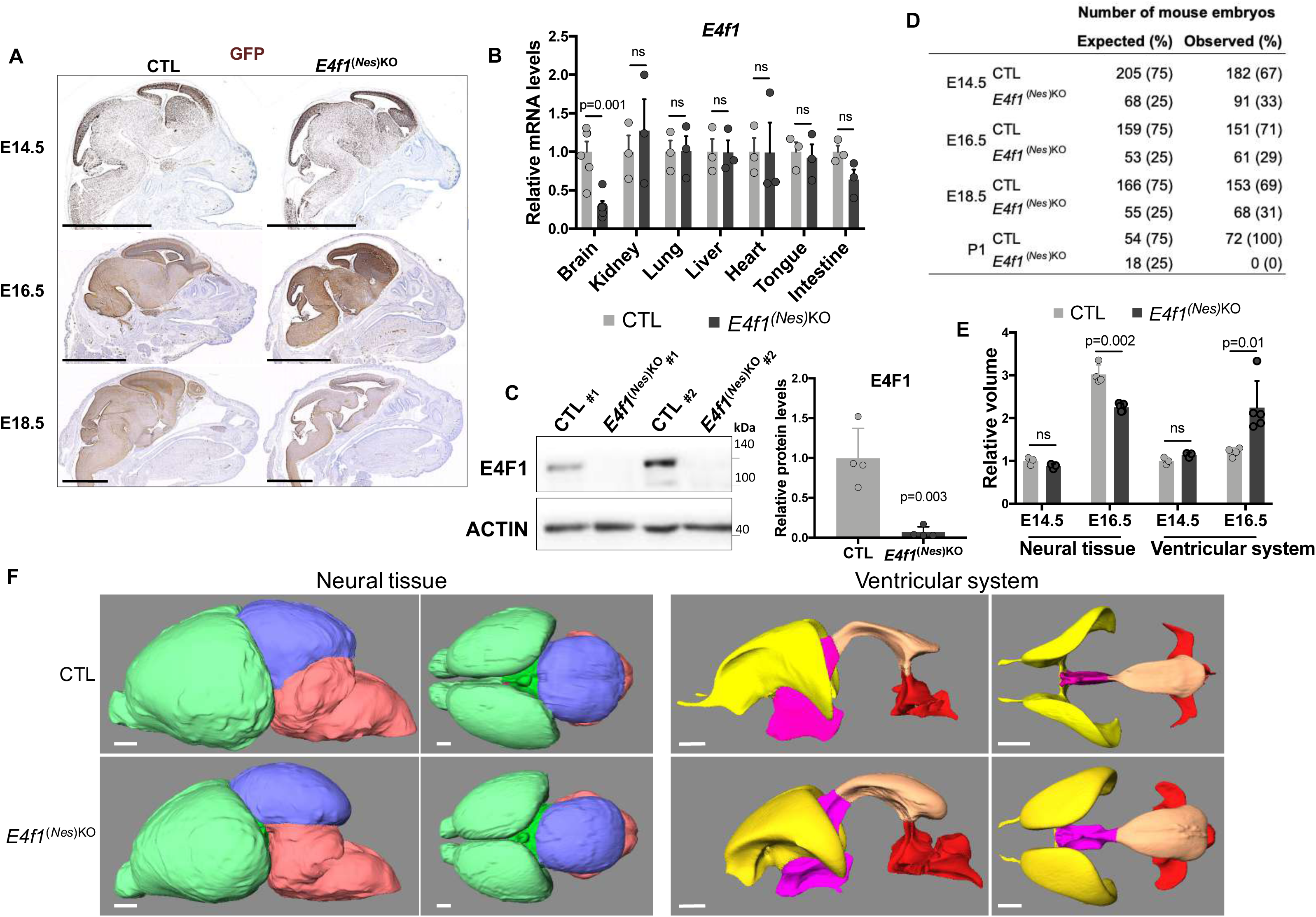
E4F1 plays a critical role in corticogenesis during mouse embryonic development. (A) Immunohistochemistry (IHC) analysis of GFP expression on brain sagittal sections prepared from E14.5, E16.5 and E18.5 *E4f1^(Nes)KO^* and CTL littermates. Scale bars, 2.5 mm. Images are representative of experiments performed on n=4 animals/group. (B) RT-qPCR analysis of *E4f1* mRNA levels in the indicated tissues in E18.5 *E4f1^(Nes)KO^* and CTL embryos (n*≥*3). (C) Immunoblot analysis of E4F1 and ACTIN protein levels in the brain of E14.5 *E4f1^(Nes)KO^* and CTL embryos. Histobars represent the quantification of n=4 immunoblots performed on independent samples for each genotype. (D) Numbers of *E4f1^(Nes)KO^* and CTL embryos identified at different developmental stages. Expected numbers were calculated based on a mendelian distribution. (E-F) Analysis of brain structures in E14.5 and E16.5 *E4f1^(Nes)KO^* embryos by High Resolution Episcopic Microscopy (HREM). (E) Histobars represent the relative volume of the neuroepithelium tissue and of brain ventricles of *E4f1^(Nes)KO^* and CTL embryos at E14.5 and E16.5 (n*≥*3). (F) Representative HREM images showing the 3D volumic reconstruction of different areas of the brain (forebrain in green, midbrain in blue and hindbrain in red) and of ventricles (lateral ventricles in yellow, third ventricle in pink, mesencephalic vesicle in orange and fourth ventricle in red) of E16.5 *E4f1^(Nes)KO^* and CTL littermates (n*≥*4). Scale bars, 280 μm. Data are presented as mean + standard error of the mean (SEM) for Fig. 1B or standard deviation (SD) for Fig. 1C and 1E from the indicated number of animals. Statistical significance was evaluated using unpaired bilateral Student’s *t*-test (ns, not significant). See also Supplemental Figure S1.

Next, we performed high-resolution episcopic microscopy (HREM) and generated three-dimension (3D) volumic brain reconstructions of E14.5 and E16.5 *E4f1^(Nes)KO^* embryos and CTL littermates to evaluate the morphology of different regions of the CNS (Fig. 1E,F and Supplemental Fig. S2A,B). These analyses revealed that the forebrain, midbrain and hindbrain neuroepithelium of E16.5 *E4f1^(Nes)KO^* embryos presented large areas of degeneration. These defects were illustrated by patchy losses of neuroepithelium or as areas of cavitation leading to alterations in the structure of all regions of the brain. The space between the brain and the cranium was also enlarged in E4F1-deficient mutants when compared to control littermates. 3D reconstructions showed that the volume of the total and subdivided neural tissues (forebrain, midbrain, hindbrain) was smaller in *E4f1^(Nes)KO^* E16.5 embryos, whereas the volume of the total and subdivided ventricles was bigger in E4F1-deficient animals than in control littermates. Minor brain lesions were detected in the forebrain, but not in the midbrain and hindbrain, at E14.5, suggesting that the brain defects resulting from E4F1 deficiency started at E14.5 and worsened at later stages of embryonic development (Fig. 1E-F and supplemental Fig. S2A,B). Histological analyses of hematoxylin and eosin (H&E)-stained sagittal sections of E14.5 and E16.5 embryos confirmed the structural lesions observed by HREM and showed that these brain defects started at E14.5 and increased at later developmental stages. The thickness of the cortical plate (CP), which is enriched in neurons, was reduced in E14.5 *E4f1^(Nes)KO^* embryos. E4F1 deficiency in the CNS resulted in decreased cellularity of all cortical layers (Supplemental Fig. S2C). No major infiltration by CD45 positive immune cells was observed at E16.5 nor at E18.5 (Supplemental Fig. S2D). Moreover, IHC and immunoblot analyses showed enhanced cleavage of the apoptotic marker caspase 3 (C-CASP3) at E16.5 and E18.5, but very mild increase at E14.5 (Fig. 2A-C). Protein levels of cleaved caspase 8, a regulator of the extrinsic apoptic pathway and of other forms of cell death, was not induced in the brain of E16.5 *E4f1^(Nes)KO^* embryos (Supplemental Fig. S2E). To identify the neural populations in which E4F1 plays an important role, we determined in coronal brain sections prepared from E14.5 embryos the number of apical and intermediate progenitors, as well as that of neurons, identified on the basis of SOX2, TBR2 and TBR1 expression, respectively. The number of TBR2 positive intermediate progenitors, and that of TBR1 neurons decreased in the brain of *E4f1^(Nes)KO^* E14.5 embryos, whereas the number of SOX2 positive apical progenitors remained comparable to that of control animals. We also evaluated the expression of C-CASP3 and of the proliferation marker Ki67 (Fig. 2D). These *in situ* analyses showed that E4F1 deficiency impaired the proportion of cycling progenitors, likely explaining the decreased cellularity observed in various regions of the neuroepithelium. Moreover, and consistently with our IHC analyses, no C-CASP3 staining was observed at E14.5, suggesting that the proliferation defects of E4F1-deficient progenitors at E14.5 preceded the neuronal cell death detected at later stages of development. These data indicate that E4F1 plays a critical role in neuronal proliferation and cell survival during mid-corticogenesis and that perturbation of E4F1 functions in the CNS results in microcephaly and perinatal death.

**Figure 2.**
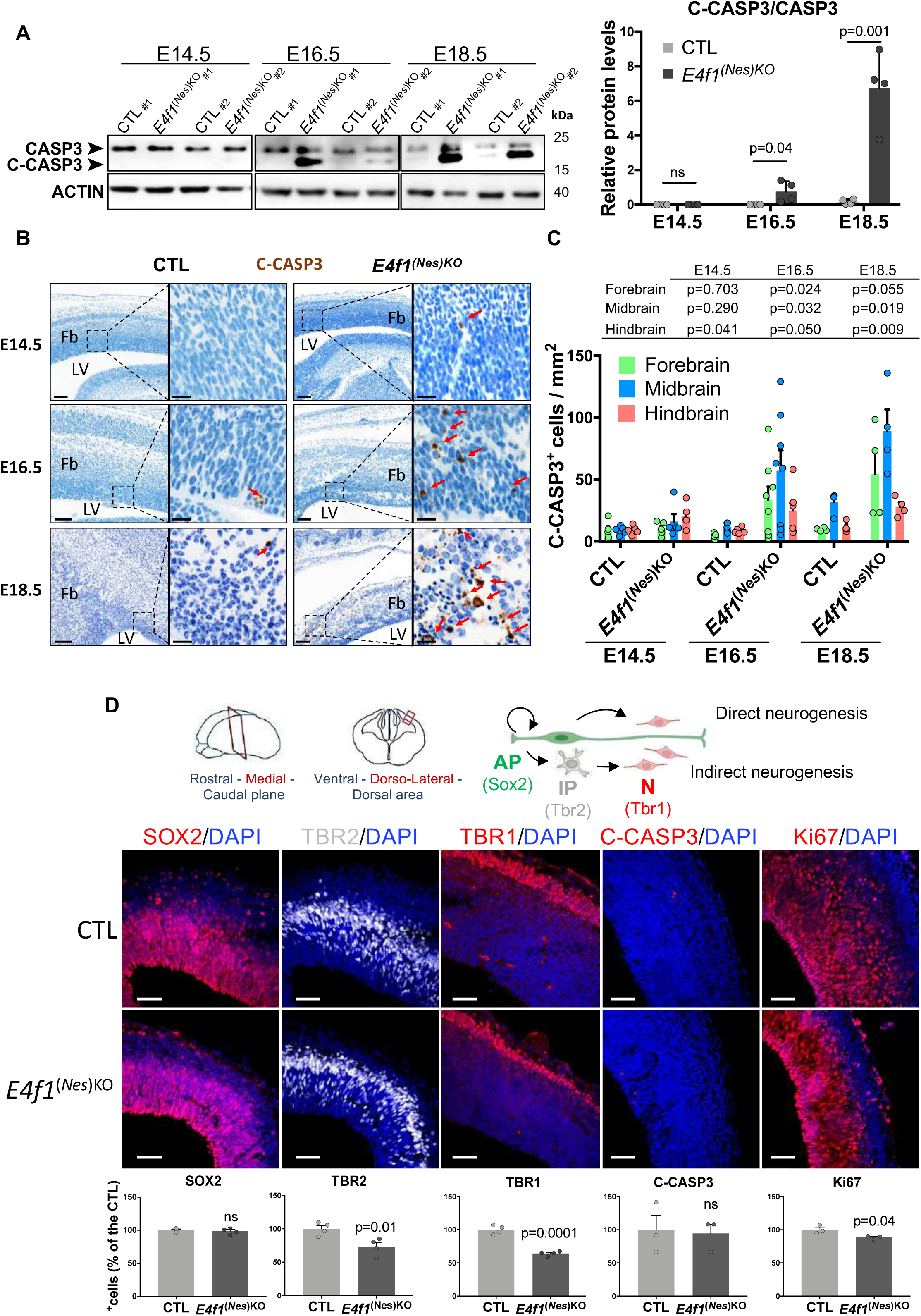
*E4f1* inactivation impairs neuronal development during mid-corticogenesis. (A) Left panels: immunoblot analysis of uncleaved and cleaved CASPASE 3 (CASP3 and C-CASP3, respectively) and ACTIN (loading control) protein levels in the brain of E14.5, E16.5 and E18.5 *E4f1^(Nes)KO^* and CTL embryos. Histobars represent the quantification of immunoblot analyses of the ratio C-CASP3/CASP3 protein levels in the brain of *E4f1^(Nes)KO^* and CTL embryos at the indicated developmental stage (n=4 animals/group). (B) IHC analysis of C-CASP3 protein in brain sagittal sections prepared from E14.5, E16.5 and E18.5 *E4f1^(Nes)KO^* and CTL embryos. Insets show images at higher magnification of the forebrain neuroepithelium. Images are representative of experiments performed on n=4 to 7 animals/group. Scale bars, 100 μm and 20 μm for the insets. Red arrows point at C-CASP3 positive cells. Fb= forebrain; LV= lateral ventricule. (C) Histobars represent the number of C-CASP3 positive cells in different brain regions of *E4f1^(Nes)KO^* and CTL embryos at the indicated developmental stage (n≥4 animals/group). (D) Upper panel: schematic representation of embryonic neurogenesis and of the dorso-lateral region of the medial plane used to determine the relative numbers of the different neuronal progenitors and their respective proliferation and survival rates by immunofluorescence. The different protein markers used to distinguish apical and intermediate progenitors and neurons are indicated. Lower panel: immunofluorescence (IF) analysis of SOX2, TBR2, TBR1, C-CASP3 and Ki67 protein levels in brain coronal sections prepared from E14.5 *E4f1^(Nes)KO^* and CTL embryos. Scale bars, 50 μm. Histobars represent the number of positive cells for the indicated protein marker expressed as % of the control (n≥3). Data are presented as mean + standard deviation (SD) for Fig. 2A or standard error of mean (SEM) for Fig. 2C and 2D from the indicated number of animals. Statistical significance was evaluated using unpaired bilateral Student’s *t*-test (ns, not significant). See also Supplemental Fig. S2

### Identification of an E4F1 core transcriptional program operating in the CNS

To gain insights into the molecular mechanisms underlying impaired neurogenesis in E4F1-deficient animals, we compared the gene expression profile of total brains harvested from *E4f1^(Nes)KO^* and CTL embryos at E14.5 using RNA sequencing (RNA-seq). Upon *E4f1* inactivation in the CNS, 185 genes displayed a fold change superior to 2 with a false discovery rate (FDR) inferior to 0.05, among which 116 were significantly down-regulated and 69 were up-regulated. Among the top 35 differentially expressed genes with the lowest false discovery rate ((FDR)-adjusted p-values) in E4F1-deficient brains were *Wdr7*, *Kif1bp*, *Neurl4*, *Fktn*, *Dlat*, *Znhit6*, *Senp8*, *Tube1*, *Elp3*, *Leo1*, *Tti2*, *Dph5*, *Taz*, *Prbm1, Ddi2*, *Mrfap1*, *Taz, Dnajc19*, *Mrpl15*, *Nsun5, Eef1g* and *Rad52* (Fig. 3A,B, Supplemental Fig. S3A and Supplemental Table S1). Interestingly, these genes were previously identified as E4F1-direct target genes in mouse fibroblasts and embryonic stem (ES) cells as well as in human breast cancer cells by chromatin immunoprecipitation (ChIP) combined with next generation sequencing (ChIP-seq)^7,8,10^. Gene set enrichment analysis (GSEA) indicated that this E4F1-signature was significantly down-regulated in the brain of *E4f1^(Nes)KO^* embryos, highlighting an E4F1 core transcriptional program operating in various cell types (Fig. 3C, Supplemental Fig. S3B and Supplemental Table S2). Impaired expression of a subset of genes composing this core E4F1 program was confirmed by RT-qPCR on mRNAs extracted from total brains or forebrains of E14.5 *E4f1^(Nes)^*^KO^ embryos (Fig. 3D and Supplemental Fig. S3C). Of note, GSEA and pathway enrichment analysis (PEA) also indicated that in addition to the deregulation of this E4F1 core program, an ISR was induced in E4F1-deficient brains (Fig. 3E-F and Supplemental Fig. S3D).

**Figure 3.**
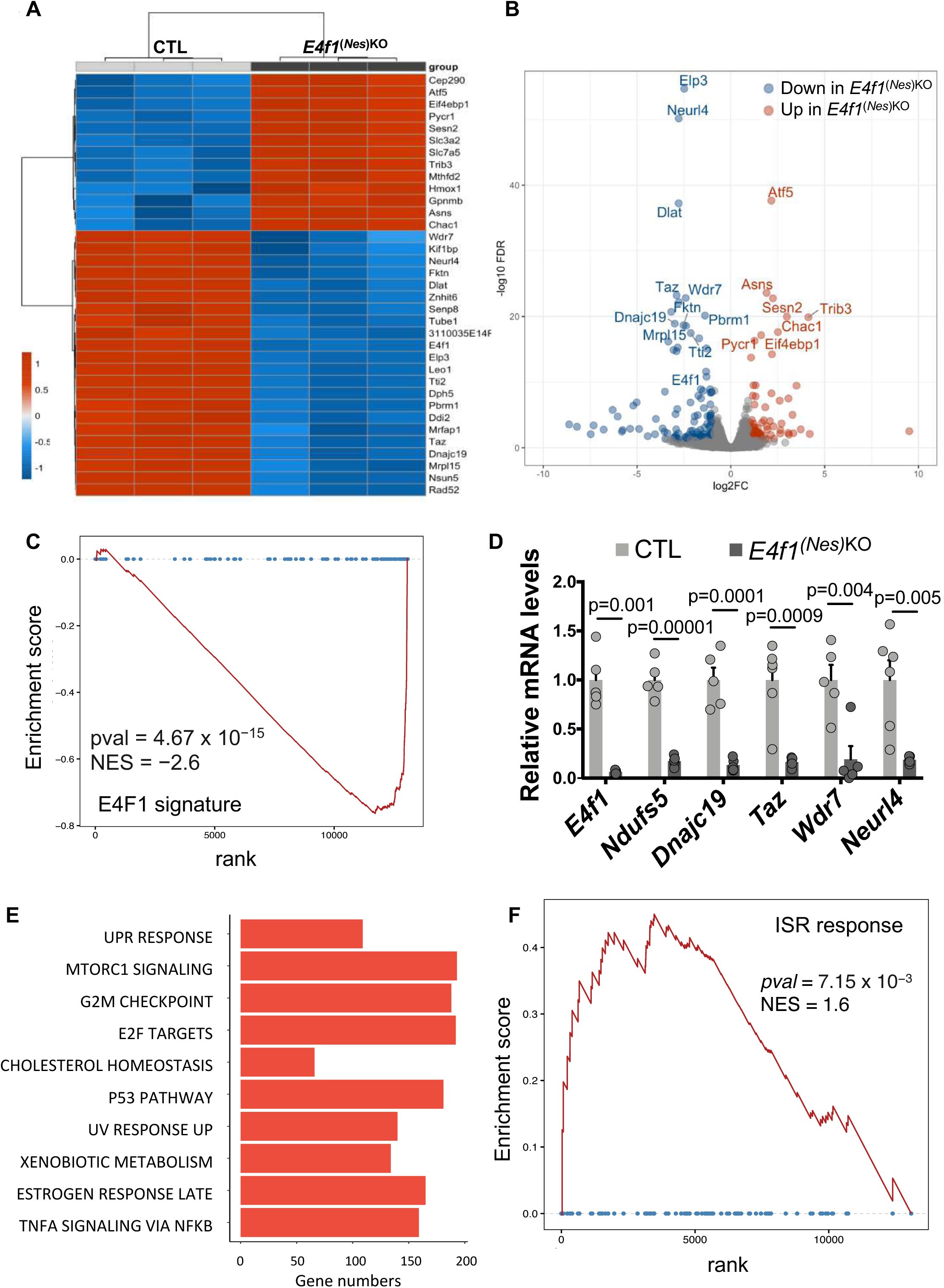
Gene expression profile of E4F1-deficient neurons. (A-B) RNA-seq analysis of brains prepared from E14.5, *E4f1^(Nes)KO^* and CTL embryos (n=3 for each genotype). (A) Heatmap representing the relative mRNA levels (fold change, log2 ratio) of the differentially expressed genes (DEG) with the most statistically significant False Discovery Rate (FDR-adjusted p-values). (B) Volcano plot depicting the significance (FDR) and magnitude of difference (fold change, log2 ratio) of DEG identified in E14.5 *E4f1^(Nes)KO^* embryos. (C) Gene set enrichment analysis (GSEA) of RNA-seq data showing decreased expression of E4F1 target genes. The normalized enrichment score (NES) and false discovery rate (FDR) are indicated. (D) RT-qPCR analysis of *E4f1* mRNA levels and those of a subset of its direct target genes (*Ndufs5*, *Dnajc19*, *Taz*, *Wdr7*, *Neurl4*) in total RNAs prepared from the forebrain of E14.5 *E4f1^(Nes)KO^* and CTL embryos (n=5). Data are presented as mean + standard error of the mean (SEM) for the indicated number of animals. Statistical significance was evaluated using unpaired bilateral Student’s *t*-test. (E) List of the top 10 pathways identified by pathway enrichment analysis (PEA) as differentially modulated in E14.5 *E4f1^(Nes)KO^* embryos. The size of the histobars represents the number of genes in the indicated category with an adjusted p value *≤* 0,001. (F) GSEA showing the induction of the Integrated Stress Response (ISR) signature in E14.5 *E4f1^(Nes)KO^* embryos. See also Supplemental Fig. S3 and Supplemental Tables S1 and S2.

### E4F1 regulates the PDC complex and pyruvate metabolism in the CNS

Among the top three most deregulated genes identified by RNA-seq in the brain of *E4f1^(Nes)KO^* embryos was *Dlat*, an E4F1-direct target gene encoding the E2 subunit of the PDC complex. Interestingly, RT-qPCR analysis of mRNAs prepared from E14.5 *E4f1^(Nes)^*^KO^ forebrains indicated that the role of E4F1 in the regulation of the PDC in neurons extended to other E4F1-direct target genes encoding several key subunits or regulators of the PDC, including *Dld*, *Mpc1*, *Pdpr* and *Slc25a19* (Fig. 4A). Furthermore, immunoblot analyses confirmed that *E4f1* inactivation in the CNS impinged on DLAT and MPC1 protein levels, whereas those of PDHA1, the E1 subunit of the PDC complex, that were evaluated as a control, as well as DLD, remained unaffected (Fig. 4B). Consistent with this expression profile, PDH activity was significantly impaired in the brain of *E4f1^(Nes)KO^* embryos and resulted in decreased amounts of AcCoA (Fig. 4C,D). Paradoxically, the abundance of histone H3 acetylation marks previously linked to nuclear PDC did not decrease in E4F1-deficient brains, suggesting that the pool of pyruvate-derived AcCoA poorly contributes to histone acetylation in the CNS (Supplemental Fig. S3E)^19–20^. Furthermore, circulating lactate levels were significantly higher in E14.5 *E4f1^(Nes)^*^KO^ embryos than in CTL animals (Fig. 4E) and IHC analyses indicated that the protein levels of the lactate transporter MCT4 was upregulated in the brain of E4F1-deficient embryos (Fig. 4F). These data show that *E4f1* inactivation in the CNS impacts on pyruvate oxidation by the PDC and favors the redirection of the glycolytic flux towards lactate production, a feature reminiscent of the metabolic reprogramming occurring in LS patients.

**Figure 4.**
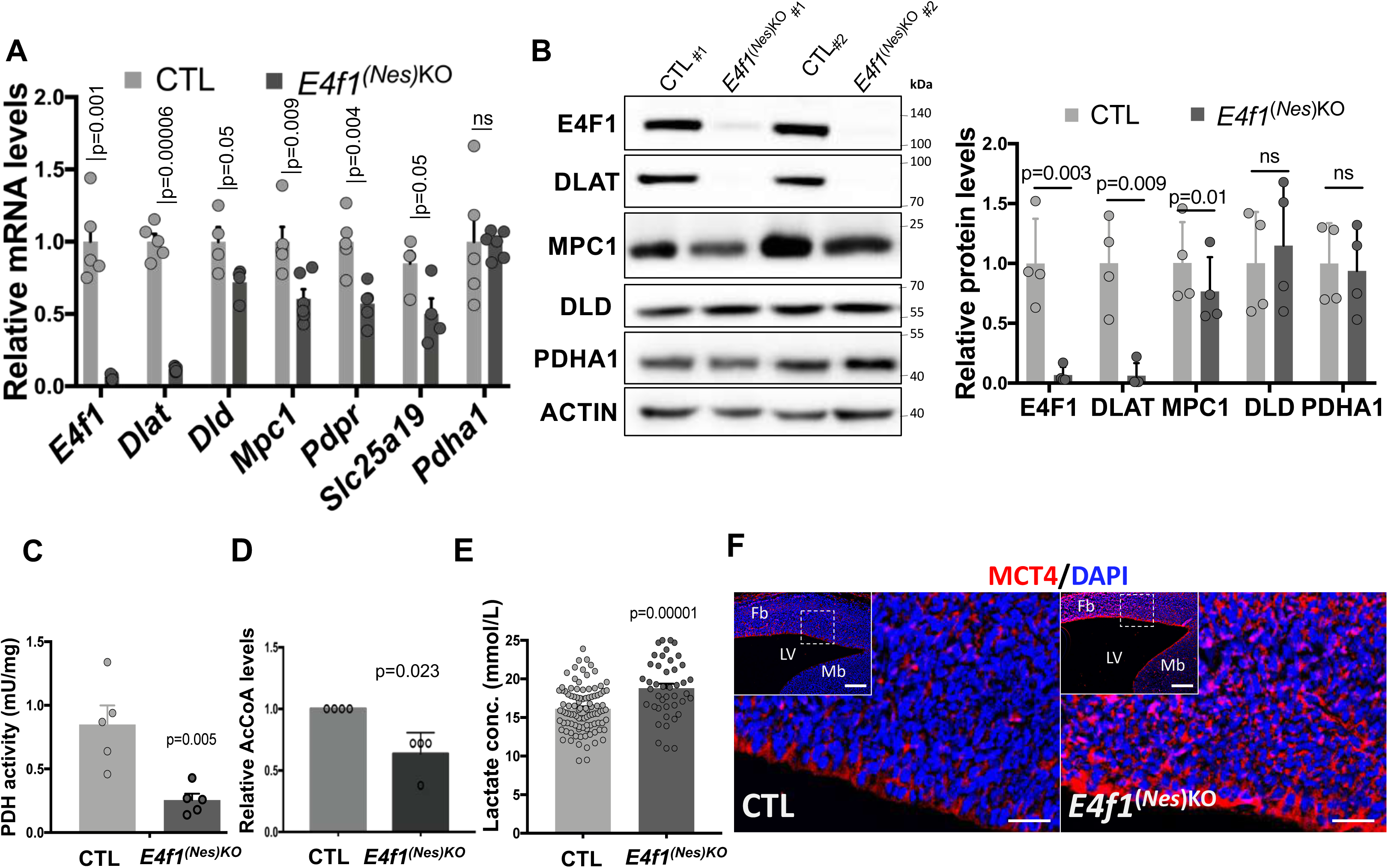
E4F1 regulates pyruvate metabolism in the CNS. (A) RT-qPCR analysis of *E4f1* mRNA levels and those of genes encoding key subunits or regulators of the pyruvate dehydrogenase complex (PDC) (*Dlat*, *Dld*, *Mpc1*, *Pdpr*, *Slc25a19* and *Pdha1*) in the forebrain of E14.5 *E4f1^(Nes)KO^* and CTL embryos (n≥4 animals/group). (B) Representative immunoblots showing E4F1, DLAT, MPC1, DLD, PDHA1, and ACTIN (loading control) protein levels in the brain of E14.5 *E4f1^(Nes)KO^* and CTL embryos. Right panel: Histobars represent the quantification of immunoblots performed on n=4 independent samples/group. (C) Pyruvate Dehydrogenase (PDH) activity in protein extracts prepared from E18.5 *E4f1^(Nes)KO^* and CTL embryos (n=5 animals/group). (D) Relative AcetylCoenzyme A (AcCoA) levels in brain from E18.5 *E4f1^(Nes)KO^* and CTL embryos measured by HR-MS (n=4 animals). (E) Blood lactate levels in E18.5 *E4f1^(Nes)KO^* and CTL embryos (n*≥*40 animals/group). (F) IF analysis of MCT4 protein levels in brain sagittal sections prepared from E14.5 *E4f1^(Nes)KO^* and CTL embryos. Fb= forebrain; Mb= middle brain; LV= lateral ventricule. Scale bars, 50 μm. Data are presented as mean + standard error of mean (SEM) for Fig. 4A and C or mean + standard deviation (SD) for Fig. 4B, D and E from the indicated number of animals. Statistical significance was evaluated using unpaired bilateral Student’s *t*-test (ns, not significant).

### *E4f1* controls the Elongator complex and U_34_ codon-biased translation in the CNS

Strikingly, our RNA-seq datasets indicated that *Elp3* mRNA levels were also strongly reduced in E4F1-deficient brains (Fig. 3A,B). *Elp3* encodes the catalytic subunit of the Elongator complex, an acetyl-transferase that was previously implicated in neurogenesis^21^. This prompted us to further investigate the links between E4F1 and the Elongator complex. Analysis of previous ChIP-seq experiments performed in murine embryonic fibroblasts (MEFs) and in ES cells suggested that *Elp3* is a direct target gene of E4F1 (Fig. 5A)^7,22^. Quantitative ChIP (qChIP) experiments performed with a previously validated polyclonal anti-E4F1 antibody^7^ confirmed that E4F1 binds to the promoter region of *Elp3* on its canonical binding site in neuronal N2A cells (Fig. 5B). We then determined by RT-qPCR the mRNA levels of different subunits of the Elongator complex in the brain of *E4f1^(Nes)^*^KO^ embryos. Consistent with the role of E4F1 in the direct regulation of *Elp3* transcription, *Elp3* mRNA levels were reduced in E14.5 E4F1-deficient brains, whereas those of *Elp1*, *Elp2* and *Elp4*, remained unchanged (Fig. 5C). At later developmental stages (E16.5 and E18.5), the mRNA levels of all these core subunits of the Elongator complex were affected by E4F1 deficiency (Supplemental Fig. S4A). Immunoblots showed that *E4f1* inactivation also impacted on ELP3 protein levels (Fig. 5D). In line with previous reports showing that the stability of the catalytic subunit ELP3 and that of the scaffold ELP1 are interdependent^21,23^, ELP1 protein levels also decreased in the brain of E14.5 *E4f1^(Nes)KO^* embryos (Fig. 5D).

**Figure 5.**
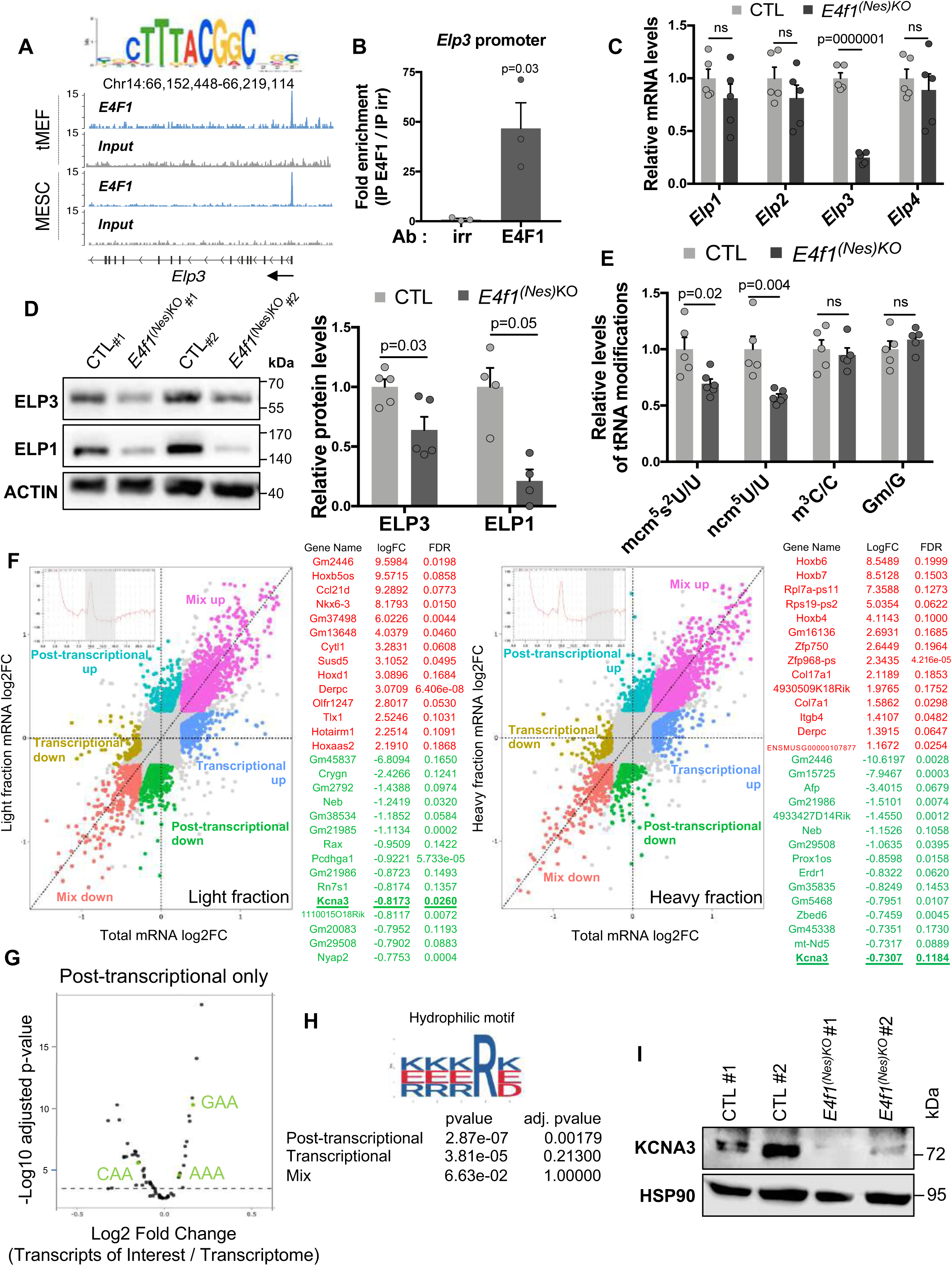
*E4f1* controls the Elongator complex and U_34_ codon-biased translation in the CNS. (A) E4F1 ChIP-seq read densities in transformed MEFs (tMEFs) and mouse ES cells (MESC) at the *Elp3* locus. The sequence and the position of the E4F1 consensus motif in the *Elp3* locus are indicated. Arrows indicate gene orientation. (B) Quantitative ChIP (qChIP) experiments on the *Elp3* promoter region using an anti-E4F1 polyclonal antibody (E4F1) or an irrelevant antibody (irr) and chromatin prepared from neuronal N2A cells (n=3 independent experiments). Results are represented as the relative ratio between the mean value of immunoprecipitated chromatin (calculated as a percentage of the input) with the anti-E4F1 antibody and the one obtained with the control antibody. (C) RT-qPCR analysis of *Elp1*, *Elp2*, *Elp3*, and *Elp4* mRNA levels in total RNAs prepared from the brain of E14.5 *E4f1^(Nes)KO^* and CTL embryos (n=5 animals/group). (D) Immunoblot analysis showing ELP1, ELP3 and ACTIN (loading control) protein levels in the brain of E14.5 *E4f1^(Nes)KO^* and CTL embryos. Right panel: histobars represent the quantification of immunoblots performed on n≥4 independent samples. (E) Liquid-chromatography coupled to tandem mass spectrometry (LC-MS/MS) analysis of the 5-carbamoylmethyl uridine (ncm^5^U_34_) and 5-methoxy-carbonyl-methyl-2-thio uridine (mcm^5^s2 U_34_) tRNAs modifications. The abundance of methyl-3-cytosine (m^3^C) and methylguanosine (Gm) that are unrelated to the Elongator complex were used as controls. The relative abundance of these tRNAs modifications was normalized to the total amount of Uridine (U), Cytosine (C) or Guanosine (G) (n=5 independent samples/group). (F) Polysome profiling analysis of brains of E14.5 *E4f1^(Nes)KO^* and CTL embryos (n=5 animals/group). Dot plots represent transcript levels (Log2 fold change) between *E4f1^(Nes)KO^* and CTL embryos in the pooled light (less then 3 ribosomes, left panel) or heavy (more than 3 ribosomes, right panel) polysomal fractions, according to their abundance in the total cytosolic fraction. The lists indicate the 16 genes which transcript abundance is the most deregulated (log^2^FC) in either the light or the heavy polysomal fractions prepared from E4F1-deficient brains but showing similar total transcript levels. (G) Codon enrichment in transcripts regulated at the post-transcriptional level in the brain of E14.5 *E4f1^(Nes)KO^* and CTL embryos. U_34_-linked codons are indicated in green. (H) Bioinformatic analysis showing that 12% of the proteins (282 out of 2288) corresponding to genes regulated at the post-transcriptional level in the brain of E14.5 *E4f1^(Nes)KO^* embryos contain at least one penta-hydrophilic amino-acid sequence previously linked to protein aggregation and degradation upon U_34_ codon-dependent translation defects. (I) Immunoblot analysis showing KCNA3 and HSP90 (loading control) protein levels in the brain of E14.5 *E4f1^(Nes)KO^* and CTL embryos. Data are presented as mean + standard error of the mean (SEM) from the indicated number of samples. Statistical significance was evaluated using unpaired bilateral Student’s *t*-test (ns, not significant). Codon enrichment was assessed by χ^2^ tests. Presence of the penta-hydrophilic sequence was determined by AME (Analysis of Motif Enrichment). See also Supplemental Fig. S4 and Supplemental Table S4.

Next, we explored the impact of E4F1-deficiency on the activity of the Elongator complex. In mammalian cells, the Elongator complex promotes the acetylation of a subset of tRNAs (tRNAs^GlnUUG^, tRNAs^GluUUC^ and tRNA^Lys^ ) on the uridine located at position 34 of the anti-codon stem loop (U_34_). This acetylation reaction initiates a stepwise cascade that first leads to the attachment of a carboxy-methyl group (cm^5^U_34_), followed by a methylation step involving the Alkylation repair homolog 8 complex (ALKBH8/TRM112) to generate 5-methoxycarbonylmethyl uridine (mcm^5^U_34_), and then the addition of a thiol group by the Cytosolic thiouridylase proteins 1/2 (CTU1/2) enzymes (mcm^5^s^2^U_34_). From ELP3-acetylated tRNAs, another branch of the pathway generates 5-carbamoylmethyl uridine (ncm^5^U_34_) through a yet unknown enzyme (Supplemental Fig. S4B)^13,14,24^. Liquid-chromatography coupled to tandem mass spectrometry (LC-MS/MS) analysis of tRNAs purified from E14.5 *E4f1^(Nes)^*^KO^ and CTL brains showed that the relative abundance of ncm^5^U_34_ and mcm^5^s^2^U_34_ decreased in E4F1-deficient neurons, whereas that of other tRNAs modifications not linked to the Elongator complex, including 3-methylcytosine (m^3^C) and methylguanosine (Gm), remained unchanged (Fig. 5E). RT-qPCR analyses showed that the total amount of tRNAs targeted by the Elongator complex, tRNAs^GlnUUG^, tRNAs^GluUUC^ and tRNA^Lys^ , were comparable in the brain of E14.5 *E4f1^(Nes)^*^KO^ and CTL embryos, demonstrating that the decreased abundance of mcm^5^s^2^U_34_ and ncm^5^U_34_ in E4F1-deficient neurons was not the consequence of an impaired production of these tRNAs (Supplemental Fig. S4C). Since the Elongator complex has also been involved in tubulin and histone acetylation, we evaluated the ratio of acetylated/total tubulin (*α*-TUB-Ac/TUB) and that of acetylated histone H3 on lysine 14/total Histone H3 (H3K14Ac/H3)^25–26^. We failed to detect any significant difference in *α*-TUB-Ac nor in H3K14Ac in the brain of E14.5 *E4f1^(Nes)^*^KO^ embryos, indicating that E4F1 deficiency in the CNS, and the associated impaired activity of the Elongator complex, preferentially impacted on tRNAs acetylation (Supplemental Fig. S4D).

Because changes in Elongator-dependent tRNAs U_34_ modifications have been linked to codon-biased translation defects, we next performed polysome profiling experiments^27^. We isolated total cytosolic mRNAs, as well as mRNAs from pooled light polysomal (1 to 3 ribosomes) and heavy polysomal (more than 3 ribosomes) fractions prepared from the brain of E14.5 *E4f1^(Nes)^*^KO^ embryos and control littermates and then determined by next generation sequencing the abundance of these transcripts (Fig. 5F and Supplemental Table S3). We first focused on a group of genes exhibiting similar total transcript levels but differential mRNA abundance in either the light or the heavy fractions purified from E4F1-deficient brains. Bioinformatic analysis of codons content showed that these mRNAs are significantly enriched in two U^34^-codons (AAA, GAA) in comparison to transcripts of the whole transcriptome (Fig. 5G). Moreover, 12% (282 out of 2288) of the corresponding proteins, including KCNA3, contain at least one penta-hydrophilic motif previously linked to protein aggregation and degradation upon codon-dependent translation defects linked to the Elongator complex (Fig. 5H)^28^. We confirmed by immunoblotting and RTqPCR that E4F1-deficiency in the brain of E14.5 embryos impacted on KCNA3 protein, but not mRNA, levels (Fig. 5I and Supplemental Fig. S4E). Altogether, these data support the notion that *E4f1* inactivation in the CNS leads to perturbed activity of the Elongator complex and translation defects that are linked, at least in part, to impaired tRNAs U_34_ modifications.

### E4F1 deficiency triggers the ISR that leads to neuronal cell death

Previous findings showed that genetic inactivation of *Elp3* during corticogenesis leads to the induction of the ISR mediated in part by ATF4, a member of the Activating Transcription Factor (ATF) family of proteins^21^. Consistent with these findings, IHC analysis of brain sections prepared from E14.5 *E4f1^(Nes)^*^KO^ embryos confirmed the induction of ATF4 protein levels (Fig. 6A). Furthermore, the induction of this ATF4-driven transcriptional response correlated with the activation of its upstream regulator eIF2a, as evidenced by its increased phosphorylation. Co-staining experiments with antibodies recognizing glial (GFAP) or neuronal (NeuN) markers indicated that both cell types exhibited increased phospho-eIF2a levels (Fig. 6B). In agreement with these data, RNA-seq and RT-qPCR analyses indicated that E4F1 deficiency in the CNS led to increased mRNA levels of ATF4-target genes implicated in the ISR, including *Atf3*, *Atf5*, *Eif4ebp1, Asns*, *Sesn2*, *Slc6a9*, *Chop*, *Chac1*, *Trib3*, *Mthfd2*, *Slc3a2* and *Slc7a5* (Figs. 3, 6C and Supplemental Table 1). The ISR involves the derepression of integral ER membrane proteins which activate independent signaling pathways^29^. Similar to *Elp3* cKO mice, the mRNA levels of *Xbp1* and its spliced transcripts *Xbp1-u* and *Xbp1-s*, as well as those of *Atf6*, remained unchanged in the brain of E14.5 *E4f1^(Nes)^*^KO^ embryos, supporting the notion that *E4f1* deficiency triggered specifically an ATF4-driven response (Fig. 6C and Supplemental Fig. S4F). Furthermore, ten days upon genetic inactivation of *E4f1* in vitro, ISR-related genes, including *Asns, Eif4ebp, Sens2, Trib3 and Slc7a3*, were also induced at the mRNA level in primary cortical cultures composed of neurons and glial cells but not in MEFs, indicating that the induction of the ISR triggered by E4F1 deficiency occurred in a cell autonomous and in a cell type specific manner (Fig. 6D and Supplemental Fig. S4G). To evaluate whether the neuronal cell death observed at E16.5 and E18.5 was linked to the induction of the ISR, we daily administered ISRIB, a selective ISR inhibitor, to pregnant females from E10.5 onwards. The induction of the ISR-related genes *Atf5*, *Eif4ebp*, *Chac1* and *Trib3* in the brain of E14.5 *E4f1^(Nes)^*^KO^ embryos was impeded by ISRIB (Fig. 6E). Furthermore, immunoblot analysis of protein extracts prepared from ISRIB or mock-treated E16.5 embryos indicated that cleaved-caspase 3 levels in the brain of E4F1-deficient embryos were reduced upon inhibition of eIF2a (Fig. 6F). However, administration of ISRIB was insufficient to rescue the neonatal lethality of *E4f1^(Nes)^*^KO^ mice (Fig. 6G). These data support a model in which perturbation of E4F1-mediated control of the Elongator complex impacts on proteome homeostasis leading to the induction of an ISR and ultimately to neuronal cell death.

**Figure 6.**
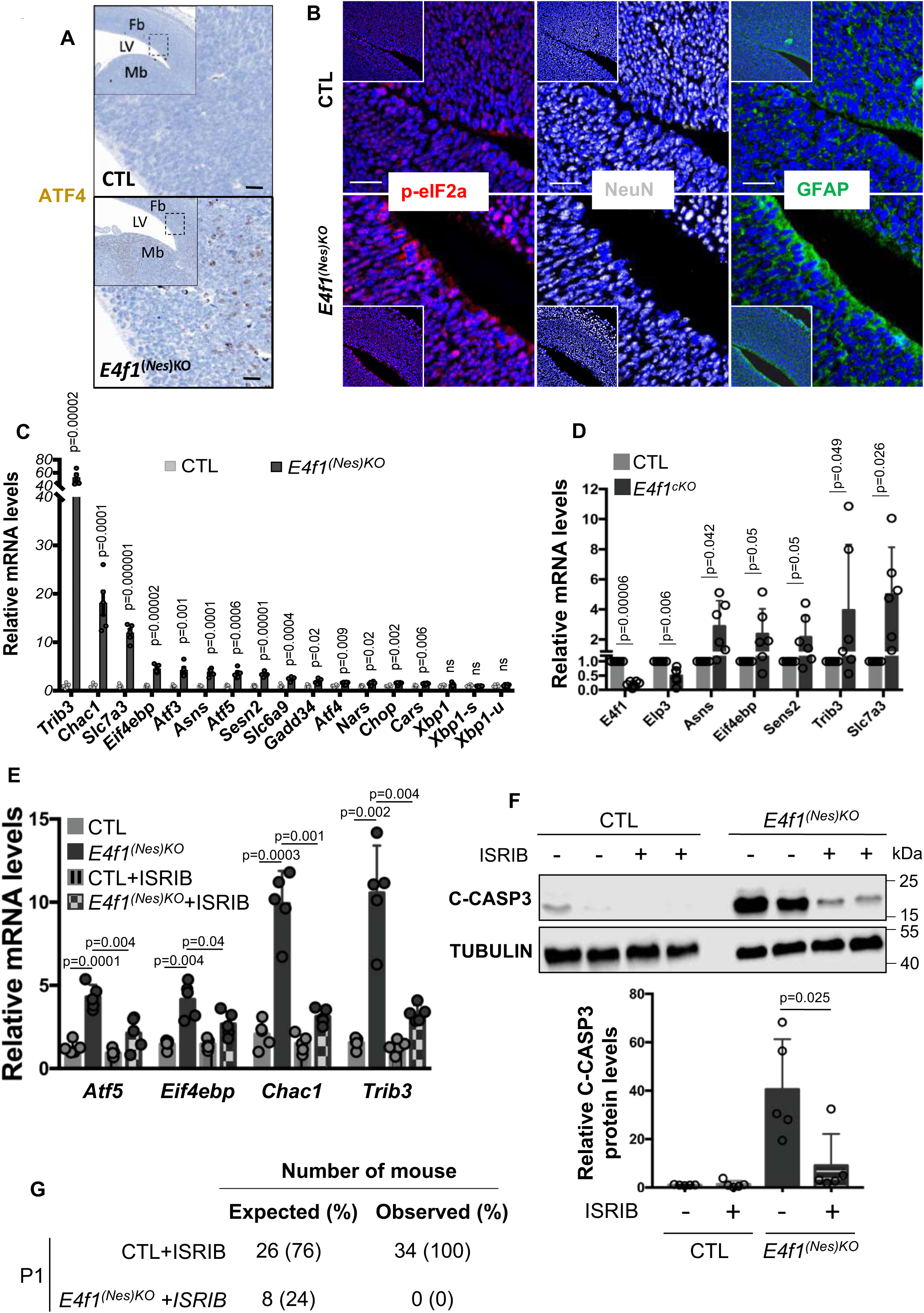
E4F1 deficiency triggers the ISR that leads to neuronal cell death. (A) IHC analysis of ATF4 protein levels in brain sagittal sections prepared from E14.5 *E4f1^(Nes)KO^* and CTL embryos. Data are representative of n=4 independent experiments. Scale bars, 100 μm. (B) Immunofluorescence (IF) analysis of phospho-eIF2a (p-eIF2a), NeuN and GFAP protein levels in brain sagittal sections prepared from E14.5 *E4f1^(Nes)KO^* and CTL embryos. Data are representative of n=4 independent experiments. Scale bars, 50 μm. (C) Relative mRNA levels of genes related to the ISR response in the forebrain of E14.5 *E4f1^(Nes)KO^* and CTL embryos determined by RT-qPCR (n=5 independent samples/group). (D) Relative mRNA levels of genes related to the ISR response in populations of *E4f1*^cKO^ and CTL primary neurons and glial cells determined by RT-qPCR, 10 days after *E4f1* inactivation in vitro (n=6 independent populations of cells/group). (E) Relative mRNA levels of genes related to the ISR response in the brain of mock or ISRIB -treated E14.5 *E4f1^(Nes)KO^* and CTL embryos determined by RT-qPCR (n=5 independent samples/group). (F) Upper panel: immunoblot analysis showing Cleaved-Caspase-3 (C-CASP3) and TUBULIN (loading control) protein levels in the brain of mock or ISRIB -treated E16.5 *E4f1^(Nes)KO^* and CTL embryos. Lower panel: histobars represent the quantification of immunoblots performed on n=5 independent samples/group. (G) Numbers of ISRIB-treated *E4f1^(Nes)KO^* and CTL embryos identified at birth (P1). Expected numbers were calculated based on a normal mendelian distribution. Data are presented as mean + standard error of the mean (SEM) for Fig. 6C or mean + standard deviation (SD) for Fig. 6D-F from the indicated number of samples. Statistical significance was evaluated using unpaired bilateral Student’s *t*-test (ns, not significant). See also Supplemental Fig. S4.

### E4F1 coordinates pyruvate-derived AcCoA production by the PDC and tRNAs U_34_ acetylation

The key role of E4F1 in pyruvate metabolism and in the regulation of the Elongator complex prompted us to evaluate whether E4F1 coordinates, at the transcriptional level, the production of pyruvate-derived AcCoA by the PDC and its utilization by the Elongator complex to promote tRNAs acetylation and ensure translation fidelity. To challenge this hypothesis, we first evaluated *Dlat* and *Elp3* expression upon acute *E4f1* inactivation in MEFs. E4F1-deficient MEFs exhibited impaired *Dlat* and *Elp3* mRNA and protein levels (Fig. 7A,B). Moreover, and consistent with our findings in *E4f1^(Nes)KO^* embryos, these cells also exhibited decreased amounts of mcm^5^s^2^U_34_ and ncm^5^U_34_ (Fig. 7C). Next, we performed stable isotope tracing experiments to evaluate whether pyruvate-derived AcCoA produced by the PDC serves as a co-factor and a carbon source for the Elongator complex to acetylate tRNAs at U_34_. MEFs were incubated in presence of ^13^C uniformally labelled pyruvate ([U-^13^C]pyruvate) for 6h and ^13^C-enrichment was measured by LC-MS/MS in the ionized fragments of ncm^5^U_34_ and mcm^5^s^2^U_34_ that were predicted to incorporate the two ^13^C-labelled carbons of the acetyl-group provided by pyruvate-derived ^13^C-AcCoA into the side chain of the uridine ring upon acetylation by the Elongator complex (Fig. 7D). The m+2 isotopologues of ncm^5^U_34_ and mcm^5^s^2^U_34_ were significantly detected in MEFs incubated with^13^C-pyruvate for 6h, a process affected upon pre-incubation with UK-5099, a pharmacological inhibitor of the pyruvate mitochondrial transporter MPC1 that indirectly blocks PDH activity (Fig. 7E and Supplemental Fig. S5A). In the same experimental set-up, ^13^C-acetate, another alternative source of AcCoA, contributed to a much lesser extent to U_34_ tRNAs modifications, suggesting that pyruvate and acetate are channeled differently in MEFs to fuel specific acetylation reactions (Supplemental Fig. S5B). These data demonstrate that pyruvate-derived AcCoA produced by the PDC is an important co-factor for Elongator-mediated acetylation reactions targeting U_34_ tRNAs.

**Figure 7.**
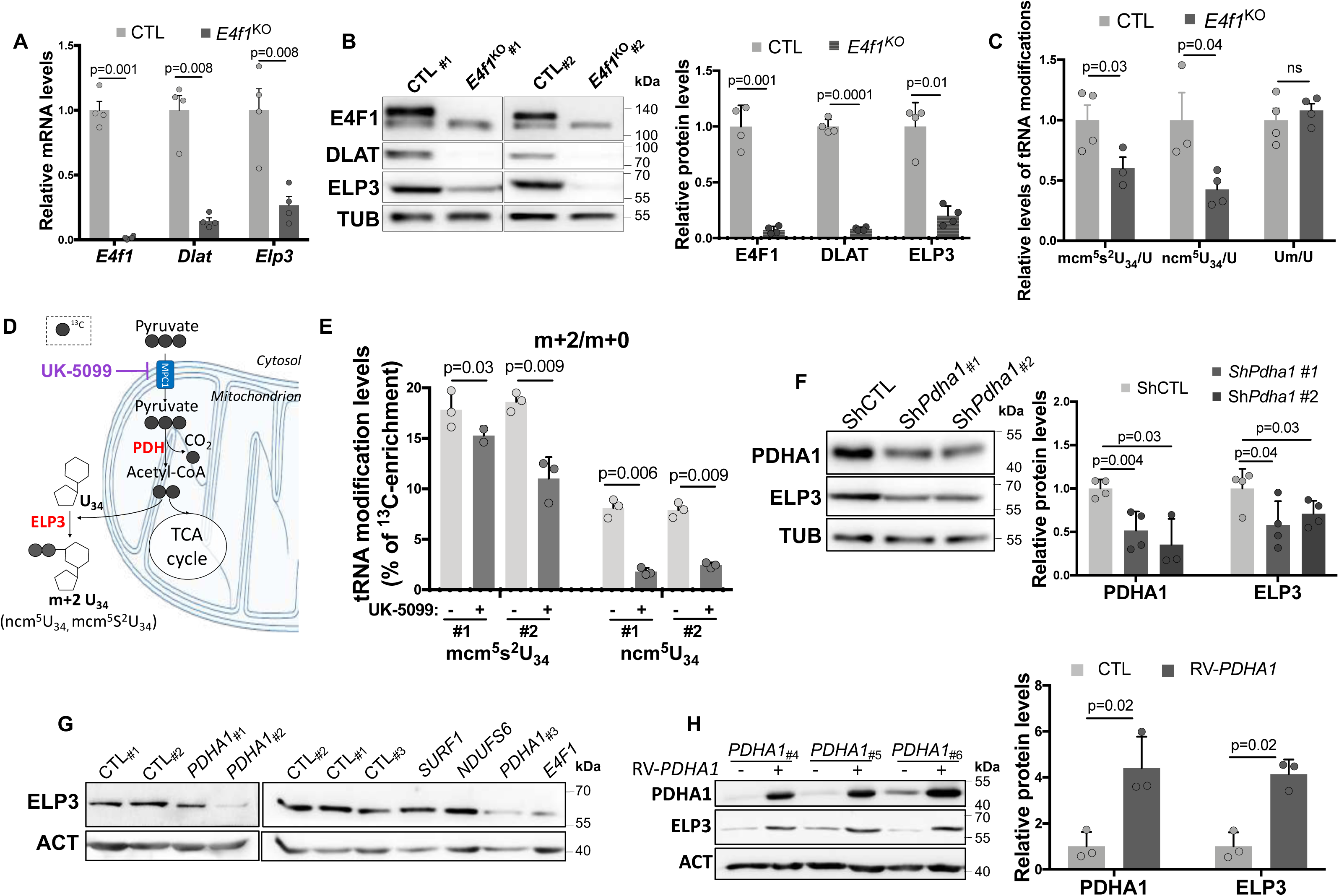
E4F1 coordinates AcCoA production by the PDC and its utilization by the Elongator complex to ensure U_34_ tRNAs acetylation. (A) RT-qPCR analysis of *E4f1*, *Dlat* and *Elp3* mRNA levels in *E4f1 cKO* and CTL MEFs (n=4 independent populations of MEFs/group), 7 days after activation of the Cre recombinase. (B) Left panel: immunoblot analysis of E4F1, DLAT, ELP3 and TUBULIN (TUB, loading control) protein levels in the same cells than in (A). Right panel: histobars represent the quantification of immunoblots performed on independent samples (n=4 populations/group). (C) LC-MS/MS analysis of ncm^5^U_34_, mcm^5^s^2^U_34_ and Um (internal control) in tRNAs purified from the same cells than in (A). The relative abundance of these tRNAs modifications was normalized to the total amount of Uridine (U) (n=4 independent populations of MEFs/group). (D) Schematic representation of the contribution of pyruvate to U_34_ tRNAs modifications when cells are incubated with uniformly labelled [U-^13^C]-pyruvate. (E) Stable isotope tracing experiments in MEFs incubated for 16 hours with DMSO or the MPC1 inhibitor UK-5099 (50 μM) and cultured for the last 6 hours in the presence of [U-^13^C]-pyruvate. ^13^C-enrichment in ncm^5^U_34_ and mcm^5^s^2^U_34_ was determined by LC-MS/MS in two independent experiments performed in triplicates. Data are represented as the relative ratio between the ^13^C-labelled m+2 isotopologue and the corresponding unlabelled U_34_ modification. (F) Immunoblot analysis of PDHA1, ELP3 and TUBULIN (loading control) protein levels in MEFs transduced with lentiviruses encoding a control or *Pdha1* shRNAs (sh*Pdha1* #1 and #2). Right panel: histobars represent the quantification of immunoblots performed on n=4 independent samples. (G) Immunoblot analysis of ELP3 and ACTIN (loading control) protein levels in human skin fibroblasts isolated from Leigh Sydrome (LS) patients harboring mutations in the indicated genes or from age-matched healthy donors. (H) Immunoblot analysis of PDHA1, ELP3 and ACTIN (loading control) protein levels in human skin fibroblasts isolated from LS patients harboring mutations in the *PDHA1* gene upon transduction with an empty control or a WT-PDHA1 encoding retrovirus. Right panel: histobars represent the quantification of immunoblots performed on cells isolated from n=3 LS patients. Data are presented as mean + standard error of the mean (SEM) for Fig. 6A and 6C or standard deviation (SD) for Fig. 6B, 6E, 6F and 6H from the indicated number of samples. Statistical significance was evaluated using unpaired bilateral Student’s *t*-test (ns, not significant). See also Supplemental Fig. S5-6.

### PDH deficiency impacts on ELP3 expression

To further understand the relationships between pyruvate metabolism and the Elongator complex, we depleted PDHA1 in MEFs using two independent short hairpin RNAs (shRNAs). Strikingly, *Pdha1* knock-down (KD) cells displayed decreased ELP3 protein, but not mRNA, levels (Fig. 7F and Supplemental Fig. S6A). Comparable decrease of ELP3 protein levels were obtained upon shRNA-mediated depletion of *Dlat* or pharmacological inhibition of PDC activity by *β*-Fluoropyruvate, unravelling a previously unknown crosstalk between the PDC and ELP3 expression (Supplemental Fig. S6B-C). Furthermore, skin fibroblasts isolated from LS patients harboring mutations in the *PDHA1* or the *E4F1* genes, but not in those with mutations in *SURF1* or *NDUFS6* displayed reduced ELP3 protein levels (Fig. 7G). Since the LS is a mendelian disease, we next evaluated the effect of ectopic wild-type (WT) PDHA1, or that of an enzymatic dead PDHA1 mutant (PDHA^C91A^), on endogenous ELP3 protein levels in these patient-derived *PDHA1*-mutated cells. Ectopic expression of WT-PDHA1, but not of PDHA1^C91A^, increased ELP3 protein levels in these *PDHA1*-mutated LS fibroblasts, confirming the notion that ELP3 expression and PDC activity are tightly coordinated (Fig. 7H and Supplemental Fig. S6D).

## DISCUSSION

The LS is a severe pediatric neurodegenerative disease linked to either PDC or ETC deficiencies. The prognosis for LS patients is very poor and current treatments provide limited improvement of their clinical symptoms. The paucity of animal models recapitulating the genetic alterations found in LS patients has limited studies aiming at understanding the molecular defects underlying their clinical symptoms and the development of efficient treatments. Moreover, because the relative contribution of cell autonomous versus systemic metabolic defects to the various clinical manifestations associated with LS remain elusive, the development of genetically engineered mouse models allowing tissue specific inactivation of genes related to the LS is of major interest. A brain specific *Pdha1*-cKO mouse model was previously generated in which PDH deficiency led to the expected metabolic reprogramming including impaired glucose-derived flux into the TCA cycle and increased lactate production. These mice also exhibited defective synthesis of neuro-transmitters linked to the TCA cycle including glutamate that was associated to neuronal excitability defects, cerebral hypotrophy and epilepsy which led to post-natal death around P25. In the same animal model, acetate and propionate were found to be efficient anaplerotic sources for the TCA which compensate, at least in part, for the lack of glucose oxidation by the PDC^30–31^. Interestingly, combining a ketogenic diet with the systemic administration of propionate mitigated the growth defects and the neuronal phenotypes of these mice but only delayed the post-natal death of these animals by a few weeks. Hence, a better understanding of the molecular defects associated to PDH deficiency is still needed to develop more efficient therapies. The detailed molecular characterization of mice in which we inactivated *E4f1*, a gene encoding a key transcriptional regulator of the PDC, in both neuronal and glial progenitors during early embryonic development uncovered an unexpected link between the PDC and the control of translation fidelity. Despite these mice display a reduction but not a complete inhibition of PDH activity, all E4F1-deficient animals died within hours after birth, likely because of reduced maternal-pup interactions and inability to feed. A comparable neonatal death was described in other mouse models developing embryonic neuronal defects, such as those resulting from the genetic inactivation of the splicing factor *Tra2b* or *Tsc1*, a component of the tuberous sclerosis complex^32–33^. Noteworthy, and consistent with our previous findings that E4F1 is a key regulator of the cell cycle and of the intra-S checkpoint^5,7,10^, E4F1-deficient animals died earlier than *Pdha1* brain-specific cKOs, strongly suggesting that E4F1 functions in the CNS extend beyond the regulation of the PDC. Using *E4f1-*brain specific cKO mice, we unravelled a network involving E4F1, the PDC and ELP3 that links pyruvate metabolism to translation fidelity. We don’t exclude the possibility that the deregulation of other E4F1-target genes, including *Nsun5* (a methyl-transferase targeting ribosomal RNAs) or *Ee1fg* (a translation initiation factor), also contributes to the translation defects observed in E4F1-deficient animals. Further investigations will be required to evaluate whether these different components of the E4F1 core transcriptional program synergistically contribute to the translation of the same subset or of different pools of mRNAs. Nevertheless, our data indicate that these mistranslated transcripts display a codon content bias towards U_34_ codons and that most of the corresponding proteins contain the hydrophilic motif previously associated with protein aggregation occurring upon perturbations of U_34_-enzymes. Altogether, these results support the notion that impaired activity of the Elongator complex plays a pivotal role in the translation defects observed in the CNS of E4F1-deficient embryos. Finally, our data also highlighted a previously unknown crosstalk between the PDC and ELP3, independent of E4F1 but relying on PDC enzymatic activity, that is perturbed in cells isolated from PDH-deficient patients, suggesting that impairement of protein translation fidelity is a common feature of cells exhibiting impaired PDH activity.

How metabolic perturbations lead to changes in gene expression is a field of active research. Many studies highlighted the role of different metabolites, including AcCoA and S-Adenosyl-Methionine (SAM), in the regulation of the epigenome. By serving as essential co-factors of acetyltransferases and methyltransferases, these metabolites directly contribute to DNA or histone modifications including acetylation or methylation, respectively^34–36^. Although it is known for decades that some of these acetyl- or methyl-transferases also target RNAs, including mRNAs, tRNAs, or ribosomal RNAs, the importance of their metabolic co-factors in regulating the epitranscriptome remains poorly understood^37^. Previous work showed that serine metabolism in mitochondria provides methyl donors to produce the taurinomethyluridine base at the wobble position of some mitochondrial tRNAs and control mitochondrial translation of OXPHOS-related transcripts encoded by the mitochondrial genome^38^. In our study, we validated an innovative stable isotope tracing approach to track the metabolic fate of pyruvate-derived carbons into U_34_ tRNAs modifications and demonstrated that AcCoA produced by the PDC directly contributes to tRNAs modifications linked to the Elongator complex. Strikingly, in the same experimental set-up, ^13^C-labelled acetate contributed much less to tRNAs U_34_ acetylation, suggesting that these metabolic substrates are channeled differently to fuel different pools of AcCoA that are utilized for specific acetylation reactions. Our data support a model where E4F1 regulates a transcriptional program coordinating AcCoA production by the PDC and its utilization by the Elongator complex to promote tRNAs U_34_ modifications. Perturbation of this program impaired protein translation fidelity and associated with the induction of an ISR that ultimately led to neuronal cell death. Interestingly, despite E4F1-mediated control of *Elp3* occurred in different cell types including MEFs, the induction of the ISR was only observed in the CNS in vivo and in neuronal cells in vitro. The mechanisms driving this cell-type specific and cell autonomous effect remain to be identified.

It is noteworthy that some antibiotics, including tetracyclines, have been shown to improve the fitness of nutrient-deprived cells isolated from patients suffering mitochondrial diseases with ETC deficiencies. Strikingly, tetracycline analogues, such as doxycycline, selectively attenuated mitochondrial translation in these cells, and when administrated to *Ndufs4*-mutant animals, improved their neuronal defects and prolonged significantly their lifespan. Although these antibiotics induced a clear ATF4-driven cellular response, their protective effects were rather attributed to the induction of a mitohormetic response which restored a proper redox balance and thereby improved cell survival^39^. These results suggest that modulating mitochondrial translation, a process amenable to pharmacological intervention, plays a pivotal role in protecting cells exhibiting ETC deficiency. In E4F1 brain-specific cKO embryos, in which we demonstrated impaired PDH activity, pharmacological inhibition of eIF2a mitigated the induction of the ISR and diminished neuronal cell death, suggesting that either the intensity or the duration of the ATF4-response may lead to different outputs in these neuronal cells. Interestingly, previous studies also suggested that inhibiting mTORC1 activity, activating a hypoxic response or modulating NAD metabolism, can partly rescue phenotypes associated to respiratory chain deficiencies^40–42^. The efficacy of these approaches in PDH-deficient-related mitochondrial diseases remains to be determined but our data pave the way for further studies aiming at evaluating whether the interplay between the PDC and the Elongator complex also represents a new avenue for therapeutic interventions for this devastating disease.

## METHODS

### Animal housing

The following strains were used in this study: *E4f1*^tm1.1Llca^, *E4f1*^tm1Pisc^, *Tg(Nes-cre)*^1Kln^, *Gt(ROSA)26Sor^tm1(cre/ERT2)Tyj^* ^15–18^. Mice were interbred and maintained on a mix 129Sv/J; C57Bl/6J background and were housed in a pathogen free barrier facility (room temperature 22 °C; relative humidity 55%, and a 12-h-light–dark cycle). All procedures were approved by the ethic committee for animal warefare of the region Languedoc Roussillon (Comité d’Ethique en Expérimentation Animal Languedoc-Roussillon), an accredited institution of the French Minister for Education, Research and Innovation (agreement number #18030-201812111250227). Animal housing and euthanasia were performed in accordance with the 3R rules. Mice were maintained under chow diet (A03, Safe) containing 22 kcal% protein, 65 kcal% carbohydrate and 13 kcal% fat. Mice were genotyped by PCR on tail genomic DNA using Red-N extract kit (Sigma), with primers indicated in Supplemental Table S4. All mouse strains are available upon request.

### Patient material

Skin fibroblasts were isolated from LS patients or aged-match control individuals recruited at Paris Kremlin Bicêtre hospital or at the Foundation IRCSS Institute of Neurology Carlo Besta (Milan, Italy). Informed consent was obtained from all investigated subjects according to the Declaration of Helsinki principles and was approved by Research Ethics committees (agreement numbers DC 2009-939 and GGP15041).

### Cell culture

MEFs were isolated from E13.5 *E4f1^-/flox^*; *Rosa26*-*CRE^ER^* embryos and cultured for no more than 3 passages in DMEM complemented with 10% of heat-inactivated fetal bovine serum (FBS, Fisher Scientific). In vitro *E4f1* inactivation in these primary cells was obtained upon incubation with 4-hydroxy-tamoxifen (1μM, Sigma) overnight (*E4f1^cKO^*). For stable isotope tracing experiments, MEFs were cultured for the last 24h in serum-free media (custom DMEM/F12 containing 5.5mM Glucose; 1mM Glutamine; 0.2mM Branched Chain Amino Acids and a mix of growth factors (ATCC, #PCS201040)), in presence of the MPC-1 inhibitor UK5099 (50μM, Sigma, PZ0160) for 12h and with 1mM of universally labelled ^13^C-pyruvate (Sigma, 490717) or ^13^C-Acetate (Sigma, 282014) for the last 6 hours. Skin fibroblasts isolated from LS patients or control individuals were cultured for no more than 8 passages in DMEM complemented with 10% of heat-inactivated fetal bovine serum (FBS, Fisher Scientific). The N2A murine neuroblastoma cell line was cultured in DMEM/F12 complemented with 10% of heat-inactivated fetal bovine serum (FBS, Fisher Scientific). Primary cortical cultures (composed of neurons and glial cells) were prepared from the brain of P1 *E4f1^flox/flox^*; *Rosa26-CRE^ER^* newborns and cultured following a previously described protocol^43^. Populations of primary skin fibroblasts isolated from LS patients and age-matched control individuals, as well as low passage populations of MEFs of interest can be provided upon request.

### RNA extraction, RNA-seq and quantitative PCR

Total RNAs were isolated from various organs or from cells using TRIzol Reagent (Invitrogen). For RNA-seq, mRNA libraries were generated following the manufacturer recommendations (mRNA hyper prep from ROCHE). Bar-coded libraries were pooled and sequenced on the Novaseq6000 200 cycles Illumina sequencer, with a sequencing depth corresponding to a minimum of 40 million of 2×100bp paired-end reads per sample after demultiplexing. Quantitative real time PCR (RT-qPCR) was performed on a LightCycler 480 (Roche). cDNAs were synthesized from 500ng of total RNA using the SuperScript™ III Reverse Transcriptase (Invitrogen). The relative mRNA copy number was calculated using Ct values and was normalized to *18S* and *tubulin β5* mRNA levels. Primers used for RT-qPCR analyses are listed in the supplemental table S3. For RNA-sequencing, STAR v2.5.3a was used to align reads on the reference genome mm10. Gene expression was determined with RSEM 1.2.28 on RefSeq catalogue, prior to normalisation and differential analysis with DESeq2 bioconductor package. Multiple hypothesis adjusted p-values were calculated with the Benjamini-Hochberg procedure to control FDR. GSEA and PEA were performed with fgsea bioconductor package on Hallmark gene sets from MSigDB collections, completed with some custom gene sets.

### Polysome profiling

Entire brain from E14.5 embryos were lysed in polysome lysis buffer (7% sucrose, 50mM Tris-HCl pH 7.5, 5 mM MgCl^2^, 25 mM KCl, 1% NP40, 10U/mL RNAseOUT supplemented with protease inhibitors and 20μg/mL Emetin) by hard shaking with 1.4mm ceramic spheres (Lysing matrix D MPBio) in a FastPrep instrument (MPBio). 30μg of total RNA were loaded on 15–50% sucrose gradient and ultracentrifuged at 210000xg for 2.5 hours at 4 °C in a SW41 rotor (Beckman Coulter). Approximately 10% of the cytosolic RNAs were kept as an input. The different fractions were separated through a live optical density (OD) 254 nm UV spectrometer and collected with an ISCO (Lincoln, NE) density gradient fractionation system. The absorbance at 254 nm was measured continuously as a function of gradient depth. Total cytosolic mRNAs, as well as mRNAs associated with 1 to 3 ribosomes (Light polysomal fraction) and mRNAs associated with more than 3 ribosomes (Heavy polysomal fraction) were extracted using TRIzol LS (Invitrogen) according to the manufacturer instructions. For sequencing, mRNA libraries were generated following the manufacturer recommendations (mRNA hyper prep from ROCHE). Bar-coded libraries were pooled and sequenced on a Novaseq6000 200 cycles Illumina sequencer, with a sequencing depth corresponding to a minimum of 80 million of 2×100bp paired-end reads per sample after demultiplexing.

Polysome profiling datasets were pre-processed using the nf-core RNA-seq pipeline v3.11.0^44^. Initial quality control of raw reads was conducted with FastQC v0.11.9. Trim Galore v0.6.7 was used for reads trimming, with parameters set to a minimum read length of 25 bp, a quality threshold of Phred score 30, and a stringency of 10. Additionally, 1 bp was clipped from both 5’ ends of R1 and R2 reads. Reads were aligned to the mouse reference genome (GRCm38) using STAR v2.7.9a. Transcript-level quantification was performed using Salmon v1.10.1, and gene-level summarization was carried out using tximport v1.32.0. Post-alignment quality was assessed with RSeQC v3.0.1 to ensure data integrity and alignment accuracy.

For statistical analysis, datasets from polysome fractions (light, heavy, and total) were analyzed independently using R v4.4.0. Differential expression (DE) analysis for each fraction was conducted with edgeR v4.2.1, comparing gene expression between *E4f1*^(Nes)KO^ and CTL animals. Normalized count data were fitted to a generalized linear model (GLM) according to *E4f1* status as a cofactor (KO or WT). After estimating the signal dispersion, the abundance of transcripts in a given fraction was considered significantly different in likelihood ratio tests when exhibiting an absolute fold-change ≥ 1.2 and a FDR-adjusted *p*-value < 0.2. To integrate results from the different fractions, the corresponding genes were categorized as controlled at the post-transcriptional level (differential abundance of their transcripts in either the heavy and/or the light polysomal fractions, but not in the total fraction), at the transcriptional level (differential abundance of their transcripts in the total fraction but not in the heavy nor in the light polysomal fractions), and at both the transcriptional and post-trancriptional (Mix) levels (differential abundance of their transcripts in the total fraction and in the heavy and/or in the light polysomal fractions). Codon usage bias was assessed by comparing codon frequencies in transcripts controlled at the post-trancriptional level in E4F1-deficient cells to those in transcripts from the entire GRCm38 transcriptome using χ^2^ tests. In the corresponding proteins, enrichment of the specific hydrophilic motif linked to U_34_-dependent translational defects was conducted using the AME (Analysis of Motif Enrichment)^45^ tool from the MEME suite v1.12.0, using the occurrence of this motif in the full GRCm38 proteome as a reference.

### Measurement of PDH activity and blood lactate levels

PDH activity was measured using Pyruvate Dehydrogenase Activity Assay Kit (MAK183-1KT, Sigma) as recommended by the manufacturer. Lactate levels were measured in blood using a lactometer (EKF Diagnostics).

### AcetylCoEnzyme A (AcCoA) quantification

Brains from E14.5 embryos were lysed in 10% trichloroacetic acid (TCA) diluted in HPLC-grade water (Fisher Scientific) precooled at 4⁰C. Samples were centrifuged and supernatants were loaded onto SPE columns (Oasis HLB 3cc; Waters WAT094226). Columns were washed with HPLC-grade water, then eluted with HPLC-grade MeOH with 25mM ammonium acetate. Samples were dried with unheated speed vac, resuspended in HPLC-grade water, transfered onto HPLC glass vials and placed in the autosampler at 4⁰C. LC–high resolution mass spectrometry quantification was performed using a Vanquish chromatographic system coupled with a Q-Exactive Plus mass spectrometer (Thermo Fisher Scientific) fitted with an electrospray source. Chromatographic separation was performed on an InfinityLab Poroshell HPH-C18 column (2.7 μm, 3.0 × 100 mm; Agilent) maintained at 30°C.

Identification was determined by extracting the accurate mass of acyl-CoAs with a mass accuracy of 5 ppm and comparing the retention time to standards injected in the same sequence (mix of 5 μM acyl-CoA). Peak areas of acyl-CoAs were divided by peak area of the internal standard used, MES, and by the weight of the corresponding brain. Data treatment was done with Xcalibur Freestyle Software (Thermo Fisher Scientific).

### Protein extraction and western blotting

Total protein extracts were prepared from total brains isolated from E14.5, E16.5 or E18.5 *E4f1^(Nes)^*^KO^ and CTL embryos lysed in Laemmli buffer (80mM Tris pH=6,8, 2% SDS, 12% sucrose, 2% β-mercaptoethanol, bromophenol blue) and immunoblotting was performed using the following antibodies: anti-E4F1 (1/1000)^46^, C-CASP3 (Cell Signaling, 9661S, 1/1000), DLAT (Santa Cruz, sc-271534, 1/500), MPC1 (Sigma, HPA045119, 1/1000), DLD (Santa Cruz, sc-365977, 1/1000), PDHE1 (Life Technologies, 459400, 1/500), ELP3 (Abcam, ab190907, 1/500), ELP1 (Abcam, ab62498, 1/500), TUBULIN (Sigma, T6199, 1/1000), ACTIN (Sigma, A3854, 1/7000), Acetylated-TUBULIN (Santa Cruz, sc23950, 1/500), Histone H3K9Ac (Cell Signaling, 9649S, 1/1000), Histone H3K23Ac (Abcam, ab61234, 1/1000), Histone H3K27Ac (Abcam, ab4729, 1/1000), Histone H3K14Ac (Cell Signaling, 7627, 1/1000), total Histone H3 (Cell Signaling, 4499S, 1/1000), HSP90 (Cell Signaling, 13998S), KCNA3 (Diagomics, AF6702, 1/500) and Cleaved-Caspase 8 (Proteintech, 66093-1).

### LC-MS detection of tRNA modifications

Total RNAs were isolated from brains of E14.5 *E4f1^(Nes)^*^KO^ and *CTL* embryos or from MEFs using TRIzol Reagent (Invitrogen). tRNAs were purified on an agarose gel. 400ng of tRNAs were digested with 0.001U of nuclease P1 (Sigma, N8630) in 25mM of ammonium acetate pH 5.3 for 2 hours at 42°C. Samples were then treated with 0.001U of alkaline phosphatase (Sigma, P4252) in 250 mM ammonium acetate pH 5.3 at 37°C for 2 hours. The nucleoside solution was diluted twice and filtered with 0.22µm filters (Millex®-GV, Millipore, SLGVR04NL). 5µL of each sample was injected in triplicate in a LC-MS/MS. For stable isotope labelling experiments performed in MEFs using ^13^C-sodium pyruvate (Sigma) or ^13^C-sodium acetate (Sigma), 4ug of total RNA were digested with 0.002 U of nuclease P1 and alkaline phosphatase by following the same procedure. Nucleosides were then separated using Nexera LC-40 systems (Shimadzu) and Synergi™ Fusion-RP C18 column (4µm particle size, 250mm x 2mm, 80Å) (Phenomenex, 00G-4424-B0). Mobile phase was composed of 5mM ammonium acetate pH5.3 (solvent A) and acetonitrile (solvent B). The 30 minutes elution gradient started with 100% phase A followed by a linear gradient until reaching 8% solvent B at 13 min. Solvent B was further increased to 40% over the next 10 minutes. After 2 minutes, solvent B was decreased back to 0% at 25.5 minutes. Initial conditions were recovered by rinsing the LC-system with 100% solvent A for additional 4.5 minutes. The flow rate was set at 0.4 mL/min and the column temperature at 35 °C. Analysis was performed using a triple quadrupole 8060NX (Shimadzu corporation, Kyoto, Japan) in positive ion mode. Peak areas were measured using Skyline 21.2 software (MacCoss Lab Software; https://skyline.ms) and the ratio of modified nucleosides on non-modified nucleosides was calculated.

### Quantitative chromatin immunoprecipitation (qChIP) assays

For qChIP assays in cells, N2A cells were incubated with 1% formaldehyde / 1% paraformaldehyde for 5 minutes. Nuclei were isolated and chromatin was extracted and sonicated (epishear, Active Motif). qChIPs on the *Elp3* locus were carried out using a control irrelevant IgG or an affinity-purified rabbit anti-E4F1 polyclonal antibody^46^, essentially as previously described^7,22^. Primers used for qChIP assays are listed in the supplemental table S4.

### Lentivirus production

Lentiviral and retroviral particles encoding Flag-tagged human WT-PDHA1, the PDHA1^C91A^ mutant, human E4F1^WT^ or the E4F1^K144Q^ mutant, and shRNAs targeting mouse *Pdha1* (TRCN0000041914, TRCN0000041914, Sigma) or mouse *Dlat* (TRCN0000041608 and TRCN0000041609, Sigma) were produced in 293T packaging cells. 72h after transfection, viral supernatants were harvested and added on MEFs overnight in presence of polybrene (5μg/mL, Sigma). Selection of transduced cells was performed 48 hours after transduction with puromycin (2μg/mL, Invitrogen) or hygromycin (50μg/mL, Invitrogen).

### Immunohistochemistry, immunofluorescence and histological analyses

The brain of E14.5, E16.5, E18.5 *E4f1^(Nes)^*^KO^ and CTL embryos were dissected in 0.1M phosphate-buffered saline pH7.4 (PBS) and were fixed at 4°C in 4% paraformaldehyde (PFA) for one hour. Fixed samples were cryoprotected overnight in 20% sucrose in PBS at 4°C, embedded in OCT Compound (VWR International) and sectioned (14µm) onto slides (SuperFrost Plus, VWR International) using a cryostat. Entire head samples were fixed in 4% neutral-buffered formalin (24h), decalcificated in 10% EDTA for 3 days and paraffin embedded. FFPE tissues were sectioned (4μm sections) and processed for immunohistochemistry (IHC) or hematoxylin and eosin (H/E) staining. For GFP (Invitrogen, A10262), Cleaved-Caspase 3 (Cell Signaling, 9661S) and CD45 (Biosciences, 14-051-82) stainings, IHC was performed on a VENTANA Discovery Ultra automated staining instrument (Ventana Medical Systems) according to the manufacturer’s instructions. The number of Cleaved-Caspase3 positive cells were quantified on whole-slides digitized sections (Panoramic 250 Flash II/3DHISTECH whole-slide digital microscope) scanned at a 0.22 µm/pixel resolution, using Definiens Developer XD software (Definiens, Munich, Germany). IF stainings were processed manually.

Briefly, brain sections were pretreated with target retrieval solution (DAKO, 20 min, 95°C) prior incubation with the following primary antibodies overnight at 4°C: anti-MCT4 (Proteintech, 22787-1-AP), ATF4 (Cell Signaling, 11815), phospho-eIF2a (Cliniscience, P04387), Ki67 (abcam, Ab15580), Ki67 (Abcam, ab15580), SOX2 (Santa Cruz, Sc17320), TBR2 (eBiosciences, 14-4875-82), TBR1 (Abcam ab31940), GFAP (Avès Labs, AB2313547) and NeuN (Biolegend, 834501). After washing, sections were incubated for 1 hour at room temperature with either anti-rabbit, anti-rat or anti-goat secondary antibodies coupled to Alexa-488, Alexa-647 or Alexa-555 (Life Technologies). Nuclei were counterstained with DAPI (Sigma) and slides were washed in PBST and coverslipped using Mowiol mounting solution. For images analyses, sections were visualized on a Thunder epifluorescent microscope (Nikon).

### High-Resolution Episcopic Microscopy (HREM)

E14.5 embryos and E16.5 heads were fixed in Bouin’s fixative for 24 hours, then washed and stored in 70 % EtOH. Fixed samples were gradually dehydrated in an increasing series of ethanol concentrations and were embedded in a methacrylate resin (JB-4 kit, Polysciences, Warrington, PA) containing eosin and acridin orange as contrasts agents as previously described^47^. The resin blocks were sectioned on the Histo3D system to generate data by repeated removal of 7 µm thick sections^48^. Resulting HREM data with a voxel size of 8 X 8 X 7 µm^3^ (E14.5 whole embryos) or 6.5 X 6.5 X 7 µm^3^ (E16.5 heads) were generated from approximately 1000 or 800 aligned images, respectively. All HREM images were converted into a volume dataset and segmented using Avizo 9.4.0 software (Thermofisher Scientific, France) to create 2- and 3-dimensional (3D) images, according to the Allen brain atlas.

### Statistical analysis

*In vivo* studies were performed on a sufficient number of animals per genotype with a minimum of 5 animals per experimental group. The in vitro data were obtained from 3 to 5 independent experiments as indicated in the figure legends. Statistical significance was evaluated using unpaired bilateral Student’s *t*-test. P values inferior or equal to 0.05 were considered statistically significant.

### Data availability

RNA-seq and polysome profiling data are available on Gene Expression Omnibus with the accessions GSE158595 & GSE278444.

## Acknowledgements

This work was supported by grants from the Agence Nationale pour la Recherche (ANR-PyrE4F), the Ligue contre le Cancer (équipe labélisée) and with the institutional support of INSERM. We are grateful for the technical support of the Montpellier Rio Imaging (MRI) and animal (RAM) facilities. Histological analyses were performed thanks to the “Réseau d’Histologie Expérimentale de Montpellier” (RHEM), a facility supported by the SIRIC Montpellier Cancer (Grant INCa_Inserm_DGOS_12553), the european regional development foundation and the Occitanie region (FEDER-FSE 2014-2020 Languedoc Roussillon), REACT-EU (Recovery Assistance for Cohesion and the Territories of Europe), IBiSA, the Ligue contre le Cancer and GIS FC3R whose funds are managed by Inserm. Implementation of the RNA mass spectrometry facility and the proteomic platform (PPM) are supported by the Occitanie region and the FEDER (PPRi and SMART projects). GA was supported by the Luxembourg National Research Fund (FNR), grant number C21/BM/15850547. We thank members of the LLC team and Massimiliano Mazzone (VIB, Leuven) for critical reading of the manuscript. MDM, MR and CDB were supported by fellowships from the laboratory of Excellence (LABEX, “Investissements d’avenir” program, ANR-10-LABX-12-01), the SIRIC Montpellier Cancer (Grant INCa-DGOS-Inserm_12553), a grant from the Institut National contre le Cancer (INCa) and the ARC foundation. We thank Y. Herault (Institut de la clinique de la souris, Illkirch) and N. Gadot and J. Valantin (pathology department at the Centre Régional contre le Cancer de Lyon), as well as J.-C. Portais (Restore, INSA, Toulouse) for their input in HREM, IHC and stable isotope tracing experiments. Next Generation Sequencing (NGS) was carried out on the iGenSeq core facility of the Institut du Cerveau (ICM) with the help of Y. Marie.

## Author Contributions

MDM, LM, NL, and LCL designed the experiments, interpreted the data and wrote the manuscript. MDM, JTC, HM, RL, TA, LS, RM, LM performed the in vivo experiments. MDM, FM, JTC, LM, PL, TI, CV, MH, AG performed in vitro experiments. GJ, CA, CP, RF and RPF designed and performed the computational analysis of RNA-seq and polysome profiling datasets. PN, BY, FPC, DP, AF, CC, ST, MHA, FFX, EB, PL, LS, WO, SGT performed the histological and HREM analyses. MDM, DBC, KBH, BF, SL, FJ, LM, AA, DA, HC designed and performed the MS and tracing experiments. LA, GD, RG, SC, LE, MDM characterized LS patient fibroblasts. All authors approved the submitted manuscript.

## Competing financial interests

The authors declare no competing financial interests.

## SUPPLEMENTAL INFORMATION

**Supplemental Figure S1.**
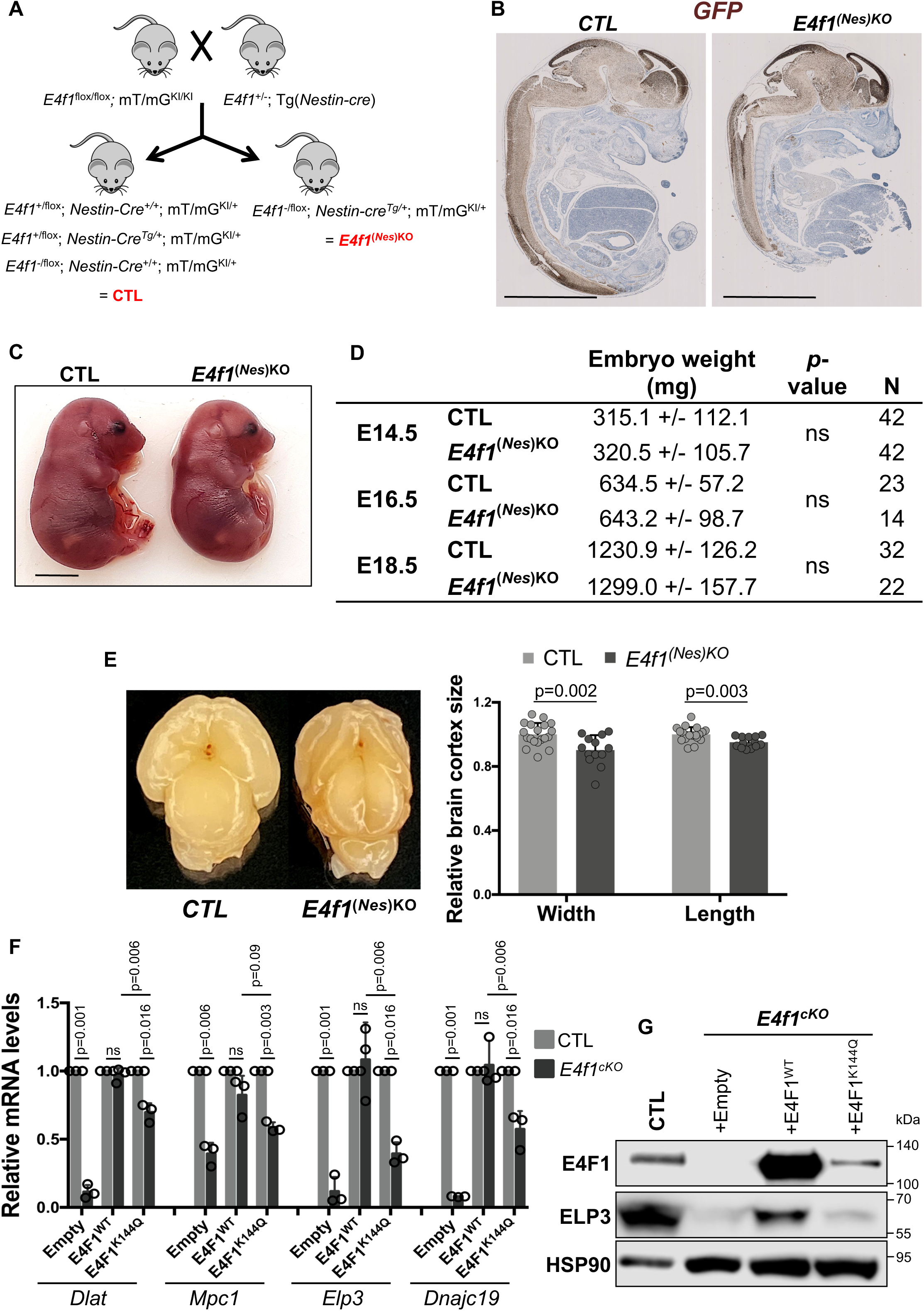
E4F1 deficiency in the CNS results in microcephaly. (A) Schematic representation of the crosses used to generate mice lacking E4F1 in the central nervous system (CNS). (B) Immunohistochemistry (IHC) analysis of GFP protein expression pattern on whole embryo sagittal sections prepared from E14.5 *E4f1^(Nes)KO^* and CTL littermates. Scale bars, 2.5 mm. Data are representative of experiments performed on n=3 animals/group. (C) Microphotograph of E18.5 *E4f1^(Nes)KO^* and CTL littermates. Scale bars, 500 μm. (D) Weight of *E4f1^(Nes)KO^* and CTL embryos at the indicated embryonic developmental stages. N indicates the number of embryos analyzed for each group. (E) Left panel: microphotograph of cortex isolated from E18.5 embryos *E4f1^(Nes)KO^* and CTL embryos. Right panel: histobars represent the width and the length of the cortex these embryos (n>13 animals/group). Data were normalized according to each embryo total body length. (F) RT-qPCR analysis of *Dlat, Mpc1*, *Elp3* and *Dnajc19* mRNA levels in *E4f1^cKO^* and CTL MEFs transduced with lentiviruses encoding E4F1 wild-type (E4F1^WT^), the E4F1 *K144Q* mutant (E4F1^K144Q^) or the empty control lentivirus (n=3 independent populations of MEFs/group). Cells were harvested 7 days after Cre-mediated inactivation of *E4f1* and transduction by the indicated lentivirus. (G) Immunoblot analysis of E4F1, ELP3 and HSP90 (loading control) protein levels in *E4f1^cKO^* and CTL MEFs transduced with lentiviruses encoding E4F1 wild type (E4F1^WT^), the E4F1 *K144Q* mutant (E4F1^K144Q^) or the empty control lentivirus. Blots are representative of 3 independent experiments. Data are presented as mean + standard deviation (SD) from the indicated number of animals. Statistical significance was evaluated using unpaired bilateral Student’s *t*-test (ns, not significant).

**Supplemental Figure S2.**
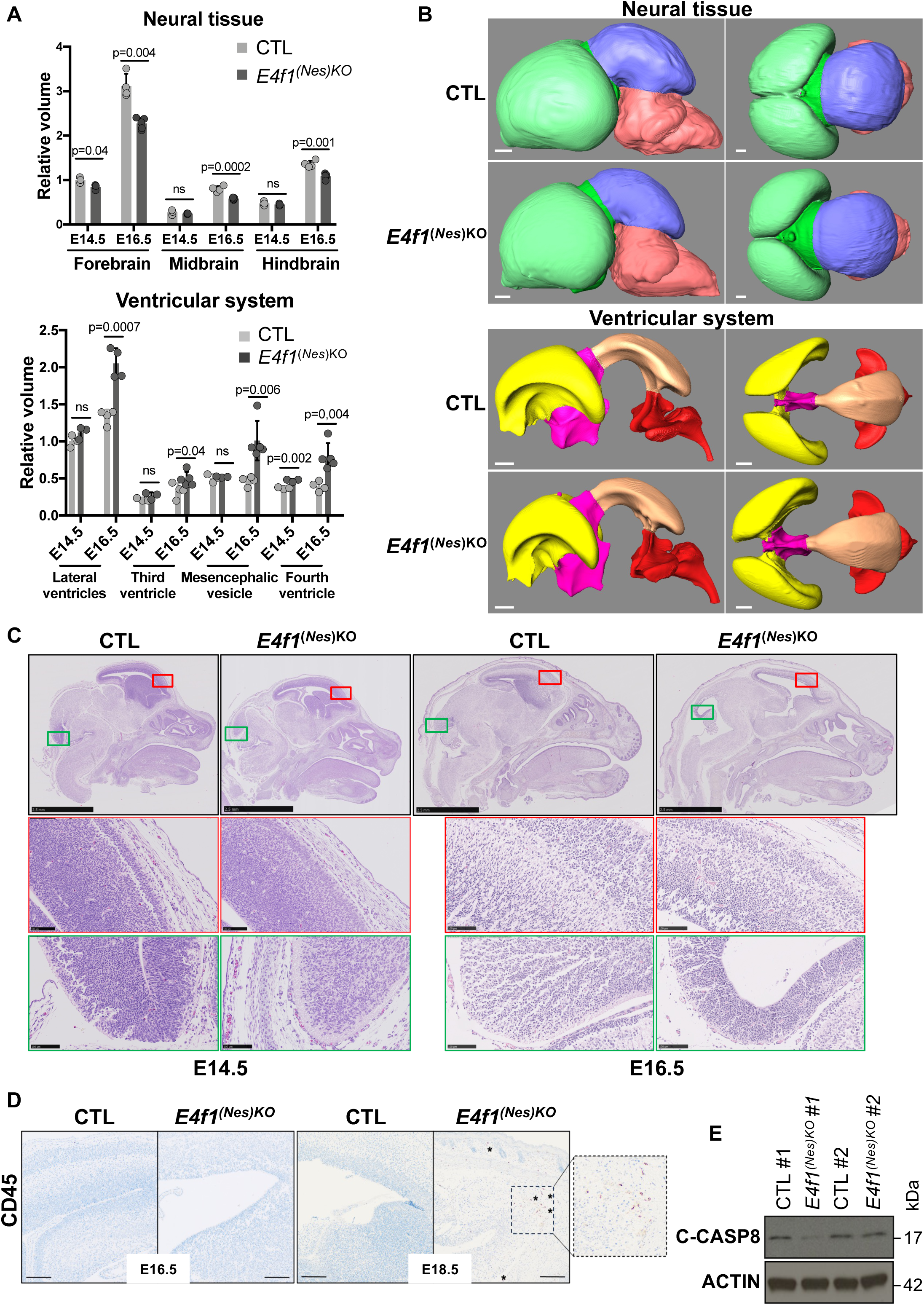
(A) Analysis of brain lesions in E14.5 and E16.5 *E4f1^(Nes)KO^* and CTL embryos by High Resolution Episcopic Microscopy (HREM). Histobars represent the relative volume of different regions of the neuroepithelium tissue and of brain ventricles of E14.5 and E16.5 *E4f1^(Nes)KO^* and CTL embryos (n≥3). Data are presented as mean + standard deviation (SD) from the indicated number of animals. Statistical significance was evaluated using unpaired bilateral Student’s *t*-test. (B) Representative HREM images showing the 3D volumic reconstruction of different areas of the brain neural tissue (forebrain in green, midbrain in blue, and hindbrain in red) and of the ventricular system (lateral ventricles in yellow, third ventricle in pink, mesencephalic vesicle in orange, fourth ventricle in red) of E14.5 *E4f1^(Nes)KO^* and CTL littermates. Scale bars, 175 μm. (C) Hematoxylin and eosin (H&E) -stained sagittal sections prepared from E14.5 and E16.5 *E4f1^(Nes)KO^* and CTL embryos. The red and green insets show images at higher magnification of the forebrain and hindbrain of the same embryos, respectively. Scale bars, 2.5 mm and 100 μm for the insets. (D) Immunohistochemistry (IHC) analysis of CD45 protein expression pattern on brain sections prepared from E16.5 and E18.5 *E4f1^(Nes)KO^* and CTL embryos. Stars indicate CD45 positive cells. Scale bars, 100 μm. (E) Immunoblot analysis of Cleaved-Caspase 8 (C-CASP8) and HSP90 (loading control) protein levels in E14.5 *E4f1^(Nes)KO^* and CTL embryos.

**Supplemental Figure S3.**
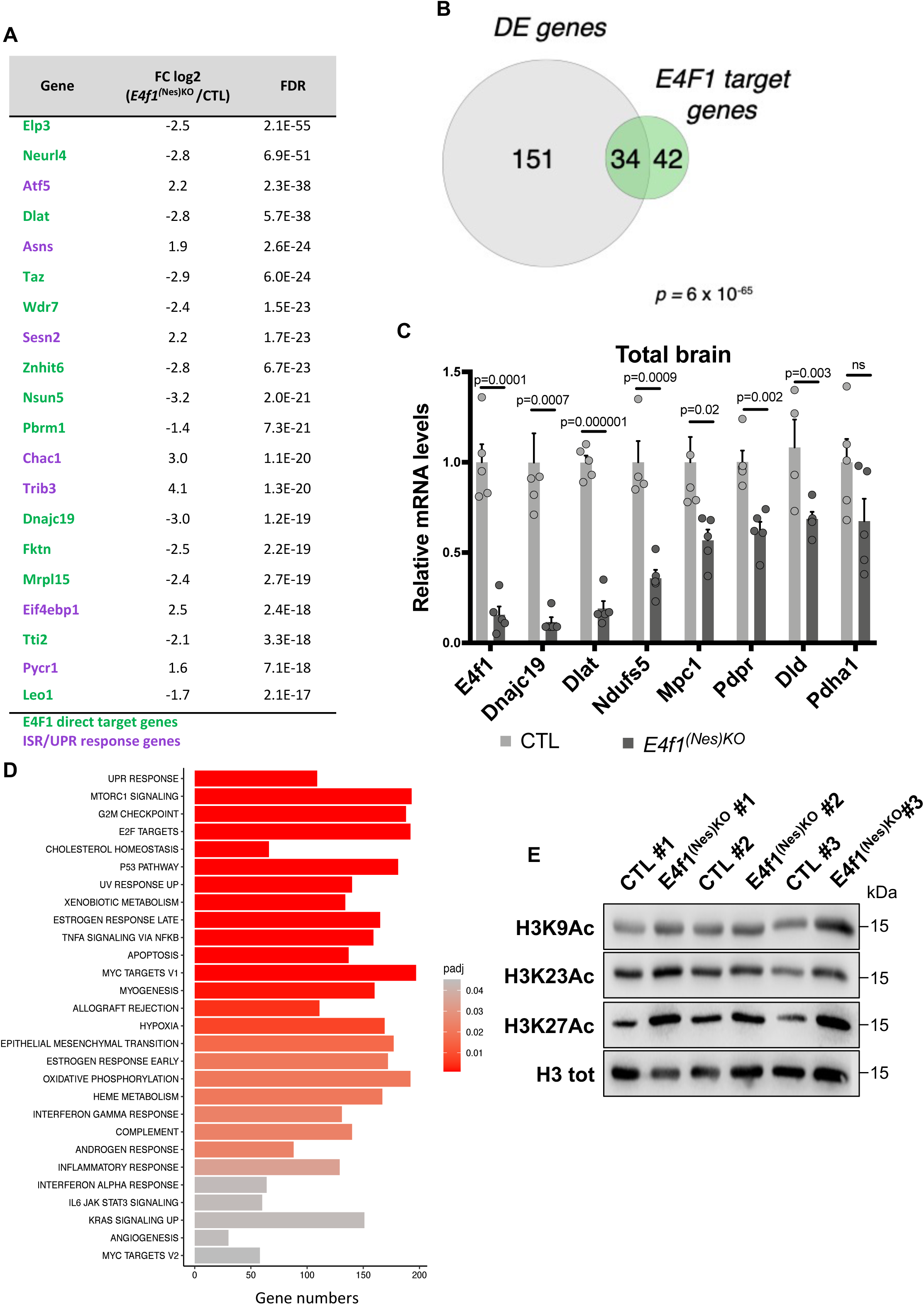
(A) RNA-seq analysis of total brain prepared from E14.5 *E4f1^(Nes)KO^* and CTL embryos. The list indicates the relative mRNA levels (fold change, log2) of the top 20 differentially expressed genes (DEG) with the most statistically significant False Discovery Rate (FDR-adjusted p-values) identified in E14.5 *E4f1^(Nes)KO^* embryos. Genes indicated in green and purple are E4F1-direct target genes and ISR-related genes, respectively. (B) Venn diagram showing the overlap between DEG identified in E14.5 *E4f1^(Nes)KO^* embryos by RNA-seq and the list of E4F1-direct target genes identified in tMEFs, MEFs, and ES cells by ChIP-seq analyses and gene expression profiling (RNA-seq and microarrays). (C) mRNA levels of *E4f1* and of a subset of its direct target genes (*Ndufs5*, *Dnajc19*, *Dlat*, *Mpc1*, *Pdpr*, and *Dld*), or of *Pdha1* (used as a control) were determined by RT-qPCR analysis using total RNAs prepared from total brain of E14.5 *E4f1^(Nes)KO^* and CTL embryos (n=5). Data are presented as mean + standard deviation (SD) from the indicated number of animals. Statistical significance was evaluated using unpaired bilateral Student’s *t*-test. (D) Pathway enrichment analysis (PEA) of DEG identified in E14.5 *E4f1^(Nes)KO^* embryos. The size of the histobars represent the number of genes in each category and the color correspond to the indicated adjusted p value. (E) Immunoblot analysis of acetylated histone H3 on lysine 9 (H3K9Ac), lysine 23 (H3K23Ac) and lysine 27 (H3K27Ac) and of total histone H3 (loading control) protein levels in E14.5 *E4f1^(Nes)KO^* and CTL embryos.

**Supplemental Figure S4.**
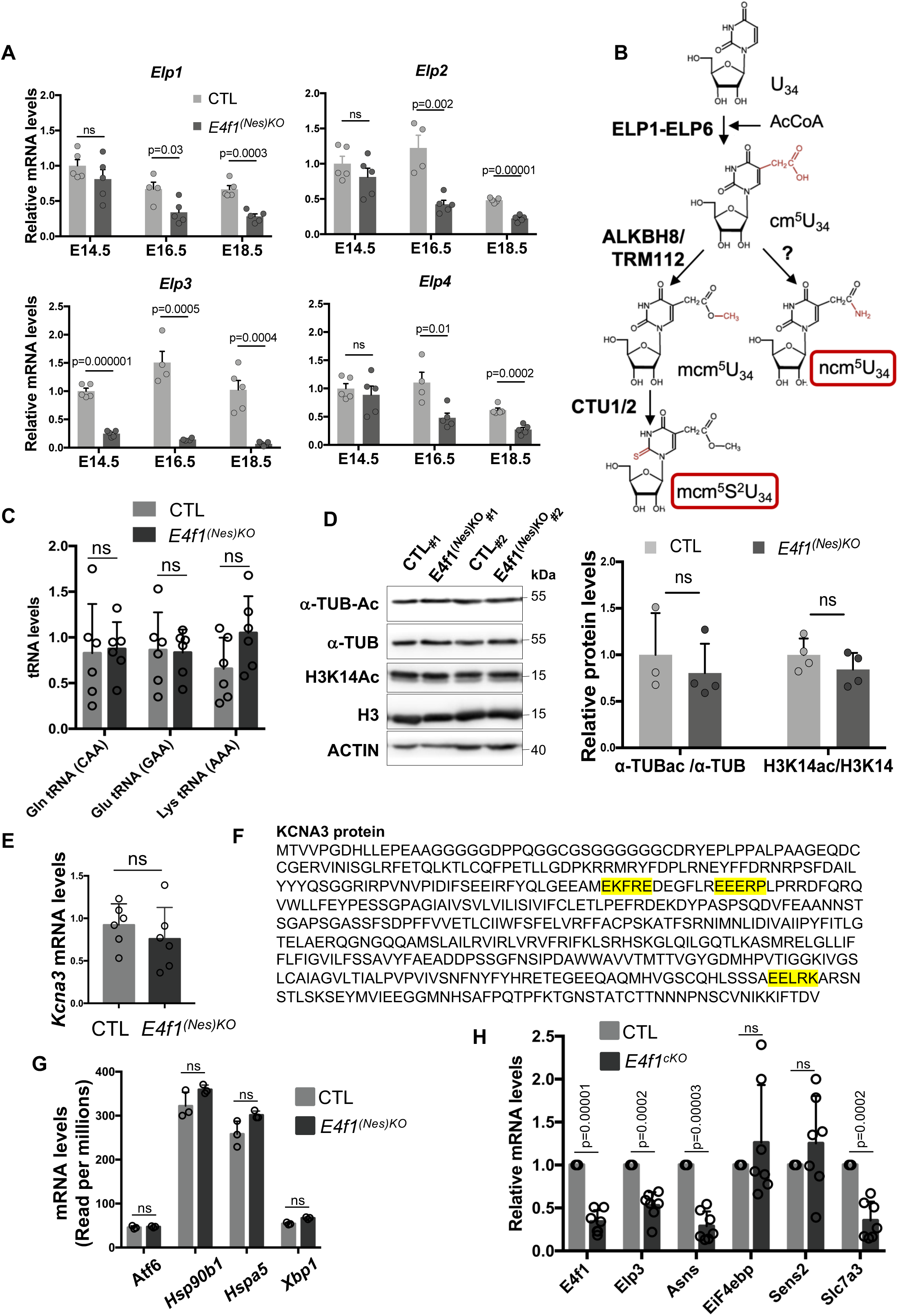
(A) RT-qPCR analysis of *Elp1*, *Elp2*, *Elp3*, *Elp4* mRNA levels in total brains of E14.5, E16.5, E18.5 *E4f1^(Nes)KO^* and CTL embryos (n=5 animals/group). (B) Schematic representation of the cascade leading to U_34_ tRNAs modifications. The enzymes catalyzing the different steps are indicated in bold. (C) RT-qPCR analysis of Gln (CAA), Glu (GAA) and Lys (AAA) tRNA levels in E14.5, *E4f1^(Nes)KO^* and CTL embryos (n=6 animals/group). (D) Immunoblot analysis of acetylated *α−*tubulin (*α−*TUB-Ac), total *α−*tubulin (*α−*TUB), acetylated lysine 14 of histone H3 (H3K14Ac), total histone H3 (H3) and ACTIN (loading control) protein levels in E14.5 *E4f1^(Nes)KO^* and CTL embryos. Right panel: histobars represent the quantification of immunoblots performed on n*≥*3 independent samples. (E) RT-qPCR analysis of *Kcna3* mRNA levels in E14.5 *E4f1^(Nes)KO^* and CTL embryos (n=6 animals/group). (F) Penta-Hydrophilic motifs linked to U_34_-codon dependent translation defects (in yellow) in KCNA3 protein sequence. (G) *Atf6, Hsp90b1, Hspa5* and *Xbp1* mRNA levels in E14.5 *E4f1^(Nes)KO^* and CTL embryos measured in the RNA-seq (Fig.3A) (n=3 animals/group). (H) Relative mRNA levels of genes related to the ISR response in *E4f1^cKO^* and CTL MEFs cells determined by RT-qPCR (n=6 independent populations of cells/group). Data are presented as mean + standard error of the mean (SEM) for Fig. S4A or mean + standard deviation (SD) for Fig. S4C-F from the indicated number of animals. Statistical significance was evaluated using unpaired bilateral Student’s *t*-test (ns, not significant).

**Supplemental Figure S5.**
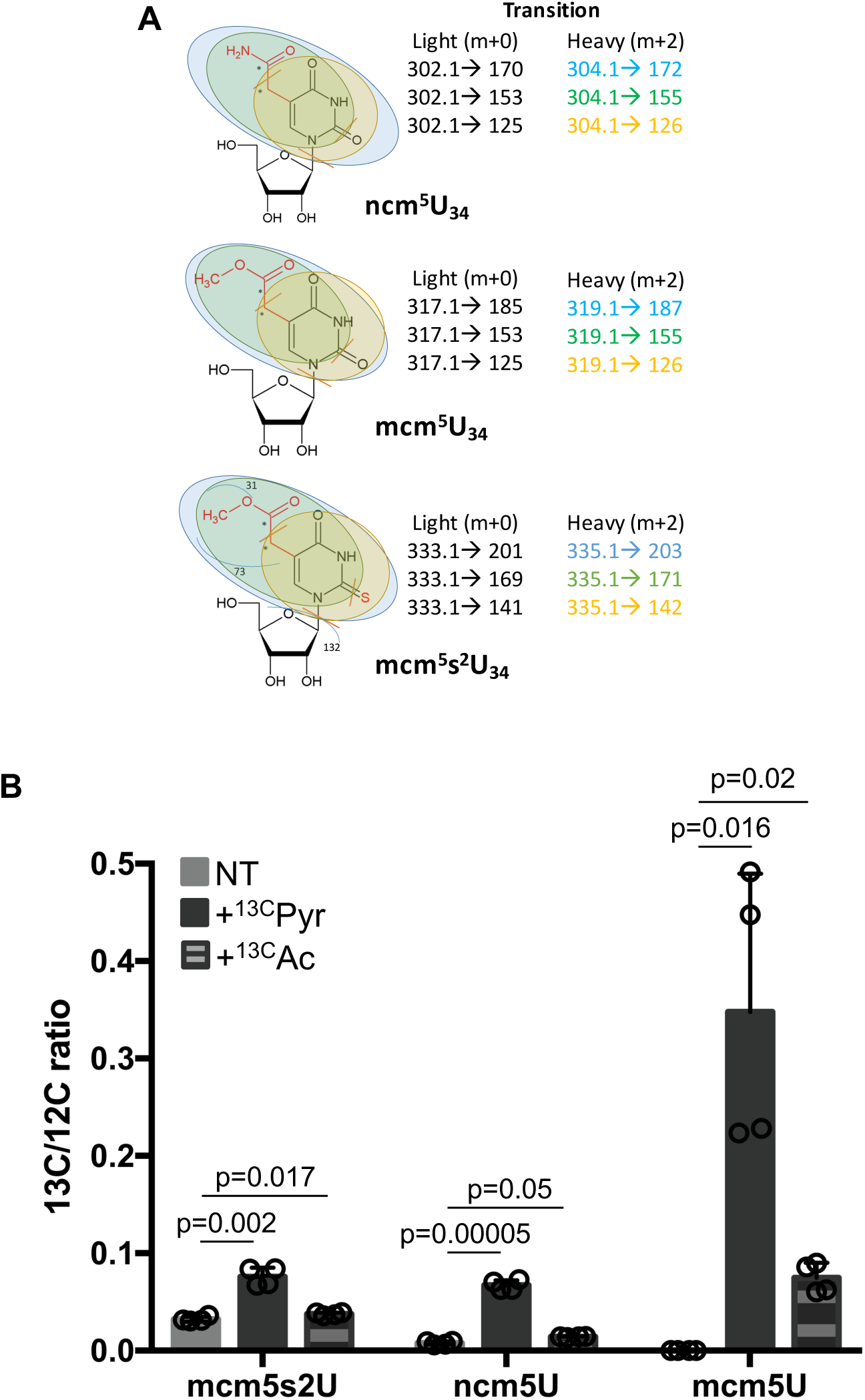
(A) Schematic representation of the different transitions and their respective molecular weight of the m+2 isotopologues of the indicated U_34_ tRNA modifications detected by LC-MS/MS upon incubation with [U-^13^C]-pyruvate. (B) Stable isotope tracing experiments in MEFs cultured for the last 6 hours in the presence of [U-^13^C]-pyruvate or acetate. ^13^C-enrichment in mcm^5^s^2^U_34_, ncm^5^U_34_ and mcm^5^U_34_ was determined by LC-MS/MS in 4 independent replicates for each condition. Data are represented as the relative ratio between the ^13^C-labelled and unlabelled U_34_ modifications. Data are presented as mean + standard deviation (SD) from the indicated number of samples. Statistical significance was evaluated using unpaired bilateral Student’s *t*-test.

**Supplemental Figure S6.**
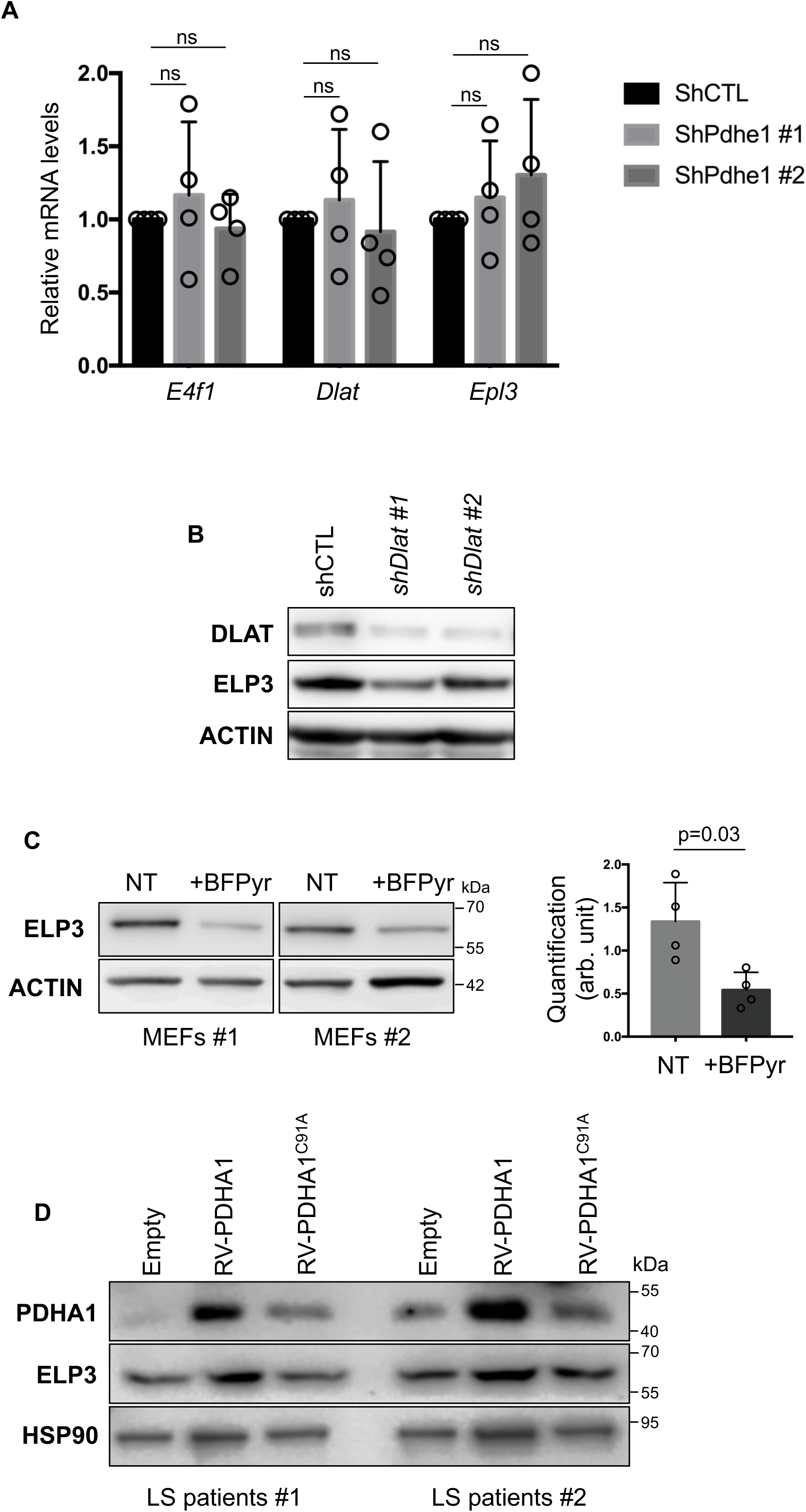
(A) RT-qPCR analysis of *E4f1*, *Dlat* and *Elp3* mRNA levels in MEFs transduced with lentiviruses encoding 2 different shRNAs targeting *Pdhe1* or a control shRNA (shCTL) (n=4 independent experiments). (B) Representative immunoblots showing EPL3 and ACTIN (loading control) protein levels in MEFs transduced with lentiviruses encoding 2 different shRNAs targeting *Dlat* or a control shRNA (shCTL). (C) Left panel: Immunoblot analysis of EPL3 and ACTIN (loading control) protein levels in MEFs treated with Beta-FluoroPyruvate (BFPyr) for 24 hrs. Right panel: histobars represent the quantification of immunoblots performed on 4 independent population of MEFs. (D) Immunoblot analysis of PDHA1, ELP3 and HSP90 (loading control) protein levels in human skin fibroblasts isolated from two LS patients harboring mutations in the *PDHA1* gene upon transduction with retroviruses encoding WT-PDHA1 (RV-PDHA1) or an activity-dead C91A PDHA1 (RV-PDHA1^C91A^) mutant or a control empty lentivirus.

**Supplemental Table S1**

Differential expression analysis based on RNA-seq data generated for the brain of E14.5 *E4f1^(Nes)KO^* vs CTL embryos.

**Supplemental Table S2**

E4F1 core transcriptional program. List of E4F1 direct target genes defined by the presence of an E4F1 ChIP-seq peak in tMEFs or mES cells, and showing differential mRNA levels between *E4f1*^cKO^ and CTL tMEFs, *E4f1^c^*^KO^ vs CTL MEFs, and between the brain of E14.5 *E4f1^(Nes)KO^* and CTL embryos.

**Supplemental Table S3**

Polysome profiling analysis of total brains prepared from E14.5 *E4f1^(Nes)KO^* and CTL embryos.

**Supplemental Table S4**

List of oligonucleotides used in this study.

## Notes

### Competing Interest Statement

The authors have declared no competing interest.

### Summary of Updates

New figures 5 and 6 New Supplemental figures

